# TriMic: a *Triticum aestivum* microbial culture collection and synthetic community for dissecting wheat-microbe interactions

**DOI:** 10.64898/2026.07.22.740170

**Authors:** Clara Tang, Athanasios Zervas, Andrea Ward, Fani Ntana, Richard L. Hahnke, Lea Ellegaard-Jensen, Peter Stougaard, Carsten Suhr Jacobsen, Amy Grunden, Manuel Kleiner

## Abstract

- Microbial culture collections and synthetic microbial communities (SynComs) are essential experimental resources for investigating underlying mechanisms of plant-microbe interactions. However, such resources remain scarce for agriculturally relevant crops. Here, we address this gap by developing TriMic, a community resource to investigate plant-microbe interactions in wheat.
- We employed a host-mediated isolation approach to obtain a bacterial collection from wheat roots and characterized its taxonomic representation and functional potential through whole-genome sequencing. We further evaluated the community dynamics of a consortium derived from this collection across time and wheat genotypes to inform the selection of a simplified community.
- TriMic includes core and keystone taxa reported in previous wheat microbiomes studies. Moreover, genomic analysis revealed genetic potential for diverse genes associated with plant colonization, nutrient cycling, plant growth promotion, and biosynthetic gene clusters with taxa exhibiting metabolic patterns associated with both generalist and specialist strategies. The proposed 15-member SynCom reflects the taxonomic and functional relevance of the full collection.
- By integrating publicly accessible bacterial isolates, whole-genome sequences, and an ecologically informed SynCom, TriMic provides a foundation for translating ecological and genomic observations into testable hypotheses, advancing mechanistic investigations of microbial community assembly and plant-microbiome interactions.

**Graphical abstract:** Overview of the workflow for developing TriMic: (a) Bacteria were isolated from wheat roots using direct and host-mediated isolation approaches (Ben Niu *et al*. 2017), followed by selection of colony morphology and time of appearance on four culture media. (b) Non-redundant collection of 88 is publicly available at the DSMZ. (c) A proposed 15-member synthetic community was developed by evaluating community dynamics of a 27-member consortium selected across the collection in wheat roots. (d) Isolate genomes were sequenced and screened for traits associated with plant-microbe interactions and secondary metabolite production.

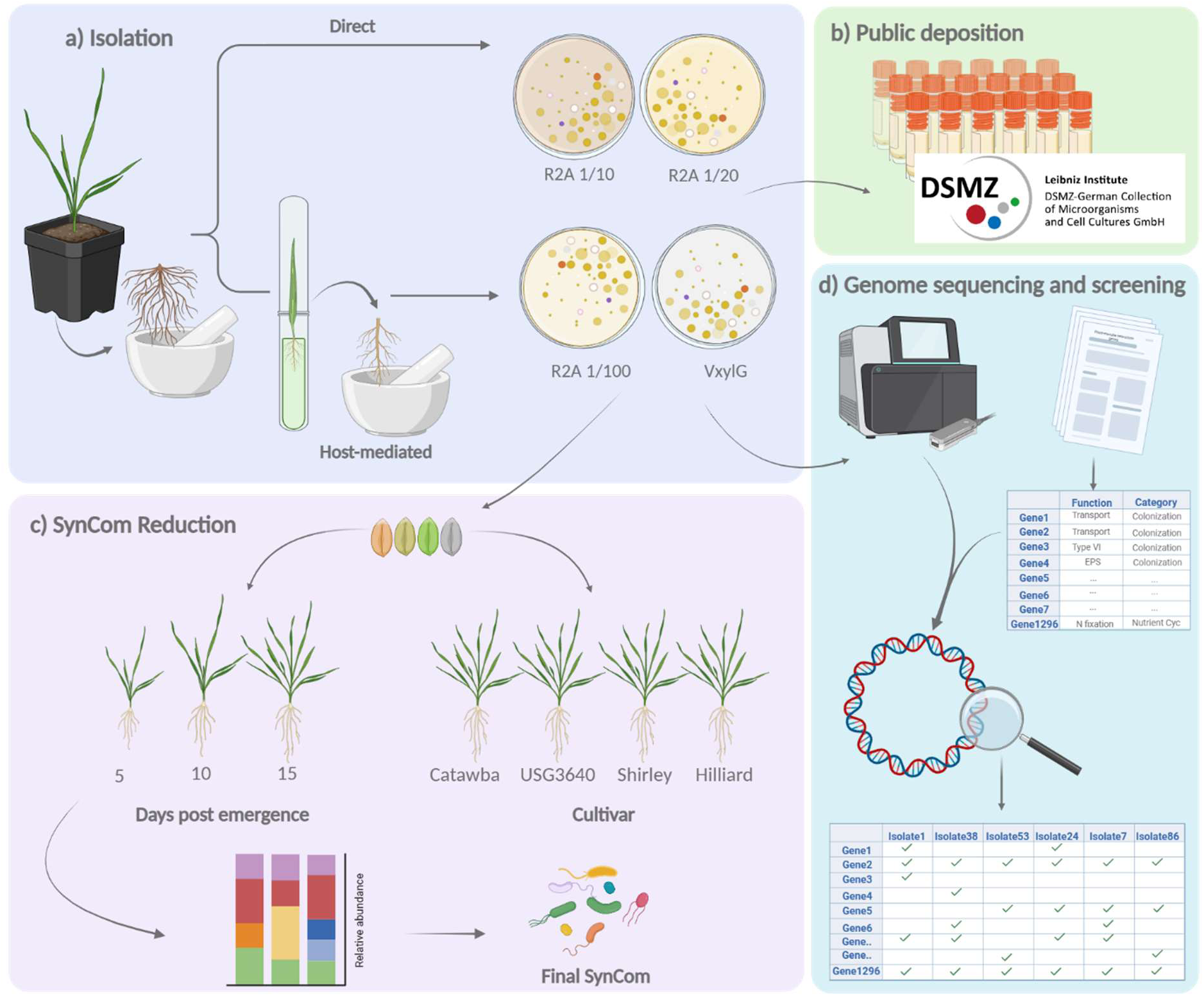

## Introduction

Managing agricultural systems to meet future global food demand calls for novel solutions that enhance crop resilience under a changing climate while minimizing the environmental cost of production. Because plant-associated microbes impact plant performance, there has been an increased interest in developing microbial products for agriculture (Batista & Singh, 2021). Microbes can enhance plant growth by improving nutrient bioavailability, increase resistance against pathogens, and improve resilience to abiotic stress (Compant *et al*., 2010; Berendsen *et al*., 2018; Trivedi *et al*., 2020). Even though microbial products (bioinoculants) have been in use for over 100 years, their widespread adoption has been low due to their inconsistent efficacy under field conditions (Batista & Singh, 2021). This inconsistent performance is partly due to a poor understanding of the modes of action of microbial beneficial traits within complex soil systems (Batista & Singh, 2021; Li *et al*., 2022). Improving the efficacy of existing and new bioinoculants will require a deep understanding of the molecular mechanisms of plant–microbe interactions (Mitter *et al*., 2021). However, the complexity of natural ecosystems hinders efforts to decipher these molecular interactions and to establish causal relationships between beneficial microbial traits and plant responses (Vorholt *et al*., 2017; Liu *et al*., 2019).

Synthetic microbial communities (SynComs) are reduced complexity systems designed to emulate natural communities (Mehlferber *et al*., 2024; Northen *et al*., 2024) and have been instrumental for discoveries in plant microbiome research. SynComs are constructed from host-derived bacterial collections that aim to capture the phylogenetic and functional diversity of the natural community, providing a foundation for mechanistic studies of plant-microbe interactions (Northen *et al*., 2024). Currently, the best characterized and most widely available plant bacterial collections are for *Arabidopsis thaliana* (Bai *et al*., 2015), *Lotus japonicus* (Wippel *et al*., 2021), and *Hordeum vulgare* (Robertson-Albertyn *et al*., 2022). SynComs derived from these culture collections, combined with ‘omics techniques, have improved experimental reproducibility and facilitated our understanding of plant host species colonization preferences (Wippel *et al*., 2021), plant organ selective colonization (Bai *et al*., 2015), microbe-microbe interactions *in planta* (Bai *et al*., 2015; Durán *et al*., 2018; Carlström *et al*., 2019), and improved plant resilience under abiotic stress (Castrillo *et al*., 2017; Levy *et al*., 2018; Rigerte *et al*., 2025). Implementing this approach to major agricultural crops to identify effector microbes or plant-growth-promoting molecules requires host-derived from globally relevant crops such as rice, wheat, soybean, and maize (Zhao *et al*., 2017).

Recently, the Crop Root Bacterial Collection was established to expand available microbial isolates to major non-model crops: maize, wheat, rice, and Medicago (Dai *et al*., 2025). Similarly, Garrell *et al*. (2026b) compiled bacterial maize isolates from different sources and sequenced their genomes to provide a highly curated and publicly available maize culture collection, which includes a SynCom containing ecologically relevant keystone taxa and fungal biocontrol capabilities developed using a host-mediated approach (Niu *et al*., 2017). These efforts represent important steps towards building a repertoire of microbial resources for agricultural crops to capture host-specific microbial traits and enable comparative studies (Busby *et al*., 2017; Northen *et al*., 2024). Wheat is important to global food security as it covers the most cultivated land area and provides one fifth of the calories and proteins consumed worldwide (Shiferaw *et al*., 2013; Erenstein *et al*., 2022). Improved understanding of the mechanisms underlying wheat-microbe interactions could facilitate the development of bioinoculants to enhance the sustainability of wheat production globally. To expand the resources available for mechanistic studies in wheat, we established a collection of root-associated bacteria using a host-mediated isolation approach in a North Carolina hard winter wheat variety. This approach leverages strong selection by the plant for the recruitment of functionally relevant microbes favoring beneficial interactions (Trivedi *et al*., 2020; Mesny *et al*., 2023). We hypothesized that: 1) The collection would contain bacterial taxa previously identified as wheat-associated microbiota and encode functions linked to plant colonization and enhanced host fitness; and 2) a SynCom derived from these taxa would retain key taxonomic members and functional attributes. Using this approach, we recovered a diverse set of bacterial isolates and curated a collection of 88 bacterial strains encompassing 31 genera, including taxa previously associated with the wheat microbiota. The collection is publicly available through the German Collection of Microorganisms and Cell cultures GmbH (DSMZ) (www.dsmz.de/collection/catalogue/microorganisms/microbiota/trimic), with bundle-priced options for the SynCom. Functional screening of the collection’s high-quality genomes revealed a broad distribution of traits associated with plant colonization, nutrient cycling, and plant growth promotion. Furthermore, we propose a SynCom derived from a consortium of isolates in the collection by tracking community dynamics in wheat roots. Together, these resources expand the experimental framework to dissect plant-microbe interactions in wheat and beyond.

## Materials and Methods

### Plant growth and root harvest

To isolate root-associated bacteria from *Triticum aestivum* L. (wheat), plants were grown at the NC State University Phytotron. Four seeds of wheat var. Catawba were grown in each D16H pot (Stuewe & Sons Inc, Tangent, USA) filled with 222 ± 0.5 mL of wheat-field agricultural soil (n=10). Pots were thinned out to a single plant after one week of growth. The soil was collected from the plough layer (0 to 25 cm) at the Cunningham Field Research Station – Lenoir site, Kingston NC. Two-hundred and eight liters of soil were quarantined in the dark at room temperature in plastic boxes for three months prior to use at the request of the phytotron to prevent introducing pests into the facility. No further handling of the soil was done prior to experiment set up. Plants were grown for 10 weeks with a 12 h photoperiod and day/night temperatures of 22/16 °C. After 2 weeks of growth, plants were vernalized for 3 weeks under an 8 h photoperiod and 4 °C temperature for day and night. Irrigation occurred every 3 days until drainage was observed from the bottom of pots. Roots were harvested using sterile equipment and cleaned by removing rhizosphere soils by shaking big aggregates off the roots and then vortexing roots for 15 s in 40 mL of sterile phosphate-buffered saline solution (PBS, Thermo Fisher Scientific, Waltham, USA) with several rinses until the solution was clear (Lakshmanan *et al*., 2017). Clean roots were cut into 0.5 cm pieces, placed in a sterile mortar with 20 mL of sterile PBS, and crushed until a slurry was obtained. Using aseptic technique, this initial root slurry was used to either inoculate surface sterilized Catawba seeds (host-mediated plating) or to directly inoculate plates (direct plating) following the two plating strategies described by Niu *et al*. (2017). A sample of the initial root slurry was stored at −20 °C for amplicon sequencing analysis of the initial community.

In the host-mediated plating strategy, wheat seeds were surface sterilized by placing them in a tea strainer and gently shaking them in the following subsequent washes: 95% ethanol for 1 min, 5.7 % available sodium hypochlorite (Clorox® Disinfecting Bleach, Oakland, USA) for 10 min, 95% ethanol for 30 s, and sterile de-ionized water for 30 s (twice) (Verma *et al*., 2019). Strainers were changed between solutions. Surface sterilized seeds were geminated for 2 days at room temperature in a sterile glass Petri dish lined with Grade 1 filter paper (Whatman 1001-125, Cytiva, UK). Once germinated, seeds were soaked in the root slurry for 1 h and sown into 0.5× Murashige & Skoog media Micro and Macronutrients (MS, RPI, Mount Prospect, USA) with 0.8% agar in a double-tube chamber (Niu *et al*., 2017). Plants were grown for two weeks at 25 °C with a photoperiod of 16 h while covering the tube containing roots with aluminum foil. Roots were harvested and a secondary root slurry was prepared as described above to inoculate plates.

### Bacterial isolation, cultivation, and identification

Isolates were obtained by plating the root slurries produced in the host-mediated and direct plating strategies. First, the root slurries were serially diluted using PBS and dilutions of 10^-2^ to 10^-5^ were plated in triplicate onto isolation media. To maximize diversity recovered, we isolated bacteria onto four different media, VXylG, 1/10 R2A, 1/20 R2A, and 1/100 R2A (Zhalnina *et al*., 2018), and colonies were selected based on their morphology and time of appearance. Plates were incubated in the dark at 25 °C for up to one week. Isolates were purified by transferring to clean plates a minimum of three times. Pure isolates were cultivated in the respective liquid media to a maximum optical density at 600 nm of 0.3. Liquid cultures were used to produce stocks, which were cryopreserved at −80 °C in 17% glycerol. Isolates were identified via polymerase chain reaction (PCR) of the full 16S rRNA gene with universal primer pair GM3F (5’-AGAGTTTGATCMTGGC-3’) and GM4R (5’-TACCTTGTTACGACTT-3’) and Taq 2X Master Mix (New England Biolabs, MA). PCR amplification was performed in 25 µL reactions in a T100 Thermal Cycler (Bio-Rad Laboratories, CA) in the following conditions: initial denaturation at 95 °C for 10 min, followed by 30 cycles of denaturation at 95 °C for 15 s, annealing at 45 °C for 60 s, extension at 68 °C for 90 s, and final extension at 68 °C for 5 min. Isolate typing was prioritized among strains with different isolation media, colony morphology, and time of appearance. All isolates were later typed. Classification, alignment, and tree inference of 16S rRNA gene sequences were done using the SILVA alignment, classification, and tree service (Pruesse *et al*., 2012). To determine the successful coverage of our collection to the original root slurry we compared our isolate identity to the community of the original root slurry. In addition, we evaluate the coverage of the collection to the known wheat microbiota by comparing our isolates to a list of wheat associated taxa compiled by Kavamura *et al*. (2021), which includes wheat microbiome studies from European, American, African, and Oceanian countries providing a non-exhaustive but clear snapshot of global wheat microbiome patterns. To focus on the most prevalent taxa while accounting for variability in the microbiome due to soil type, location, and management practices across studies (Dastogeer *et al*., 2020), we further classified this list into taxa of high prevalence (i.e. taxa shared in at least 40% of studies included), core taxa (i.e. taxa shared in at least 60% of studies included), and low prevalence (i.e. taxa shared in less than 40% of studies included). To enable this comparison, we grouped closely related genera that might not be differentiated with amplicon sequencing (*Priestia, Peribacillus*, and *Gottfrieda* were grouped into *Bacillus, Rugamonas* was designated within *Duganella*, and *Telluria* was presented as part of the *Massilia* genus).

### Whole genome sequencing

Isolates were randomly selected from each clade of the 16S rRNA phylogenetic tree for whole genome sequencing (n=100), with the number of isolates per genera representing the number of isolates recovered in each genus. Bacterial biomass for DNA extraction was produced by growing the isolates in five mL of tryptic soy broth (TSB) without dextrose (BD, NJ) at 25 °C shaking at 165 rpm until an OD_600_ was at least 0.3. Aliquots of one mL of liquid cultures were pelleted at 8000 x g for 7 min at 4 °C. DNA for short-read sequencing was extracted using the QIAGEN® Powersoil extraction kit (Qiagen, Hilden, Germany). Illumina libraries were prepared using the Nextera(R) XT DNA library Preparation kit (Illumina, San Diego, USA). Libraries were pooled equimolarly and sequenced on a NextSeq 500 (Illumina Inc, San Diego, USA) as paired-end reads using the NextSeq 500/550 Mid Output kit V2 (2 x 150 bp) flow cell. Raw read quality trimming and filtering was performed using TrimGalore (Krueger, 2021). Short read assemblies were produced using SPAdes v3.13.1 (Bankevich *et al*., 2012). Short read assemblies were evaluated using CheckM (Parks *et al*., 2015) v.1.0.7 and BUSCO (Seppey *et al*., 2019) v5.2.2. Genomes were taxonomically classified using the classify workflow in GTDB-tk (Chaumeil *et al*., 2020) v.2.4.0 (Ref v. 220). Representative genomes were selected for long read sequencing (n=65) using the dereplicate function in dRep v2.2.3 (Olm *et al*., 2017) by selecting the most complete genome in clusters with average nucleotide identity (ANI) above 99% from each MASH cluster (Supporting Information Fig S1). DNA for long-read sequencing was obtained from four mL of culture harvested as described above using the MasterPure Complete DNA and RNA Purification Kit (LGC Biosearch Technologies, Hoddesdon, UK). Nanopore libraries were prepared using the Rapid Barcoding Kit 96 (SQK-RBK110.96), Oxford Nanopore Technologies, Oxford, UK) and subsequently sequenced on a SpotON Flow cell R9.4.1 using a MinION sequencer (FLO-MIN109), Oxford Nanopore Technologies, Oxford, UK). Long read assemblies were produced with Flye(Kolmogorov *et al*., 2019) v2.9 and error corrected using NextPolish (Hu *et al*., 2020) v1.4. Isolates CT11-137 and CT11-29 were assembled using Unicycler (Wick *et al*., 2017) v0.4.7. Hybrid assemblies were evaluated and taxonomically classified as described above. Completeness of all assemblies was assessed via visual evaluation of assembly graphs in Bandage (Wick *et al*., 2015) v0.9.0 and closest typed strains were designated using the Type (Strain) Genome Server (TYGS) (Meier-Kolthoff & Göker, 2019). All phylogenetic trees were constructed from a 120 single-copy universal bacterial marker gene set identified using GTDB-tk. The tree was inferred using FastTree v2.1 (Price *et al*., 2010) under the WAG substitution model with Shimodaira-Hasegawa-like support values. The tree branch length scale represents the number of substitutions per site. Trees were visualized and annotated using the Interactive Tree Of Life (iTOL) v6 online tool (Letunic & Bork, 2024). Genome sequencing data is available from NCBI archives BioProject PRJNA1147241. To remove potentially redundant strains prior to deposition at the DSMZ while including potential single nucleotide variants, we identified single nucleotide polymorphism (SNP) using Snippy (Seemann, 2015) v4.6.0 on strains sharing above 99% ANI of their genomes. Isolates of potentially redundant strains with at least one SNP within the coding region of a gene that led to a missense, frameshift or stop mutation were both included within the collection deposition.

### Genome screening for plant adaptation and beneficial functions

To determine the genomic functional potential of isolates in the collection, a list of target Kyoto Encyclopedia of Genes and Genomes (KEGG) (Kanehisa *et al*., 2016) ortholog numbers (KO) was compiled for bacterial genomic features associated with adaptations to plants (Levy *et al*., 2018), plant and root colonization (Cole *et al*., 2017; Vannier *et al*., 2023; Knights *et al*., 2024; Selten *et al*., 2026), rhizosphere persistence (Knights *et al*., 2024), plant growth promotion (Bruto *et al*., 2014), and N and P cycling (Tu *et al*., 2019; Zeng *et al*., 2022). Functional annotation were retrieved from the KEGG database. Genes were then categorized into 93 functions within 27 functional categories using the KOs BRITE Hierarchy annotations or nutrient cycling and plant growth promotion categories (Supporting Data Table S1). Genes lacking specific KEGG functional annotations were retained based on supporting evidence from the cited literature. Genomes were annotated with eggNOG mapper(Cantalapiedra *et al*., 2021) v2.1.7. KEGG ortholog numbers were assigned to predicted protein-coding genes using Diamond (Buchfink *et al*., 2021) with a percent amino acid sequence identity of 70% and e-value of 0.001. The genomes were then screened for the presence of target KOs. The proportion of genes found in select categorical functions was calculated by dividing the number of KOs identified in the genome by the total number of KOs in that category. In addition, biosynthetic gene clusters (BGC) for secondary metabolite production were identified using antiSMASH v6.0 (Medema *et al*., 2011).

### Consortium preparation, wheat growth under gnotobiotic conditions, and plant harvest

A consortium representative of the isolate collection was constructed based on the 16S rRNA gene phylogeny. Within each genus, phylogenetic clusters were visually identified and representative isolates were randomly selected. This approach resulted in 27 isolates representing 24 genera within the collection. At the time of selecting isolates for the consortium, we had classified the collection isolates into 24 genera based on 16S rRNA sequences, however, whole genome sequencing later resolved the collection into 31 genera, thus not all 31 genera were included in the consortium.

All strains of the consortium were inoculated from cryopreserved stocks onto R2A plates and a single colony was transferred to a starter culture of five mL of TSB without dextrose prior to transferring to the final 30 mL TSB culture following the schedule and media strength in Table 1. Media strength was selected based on best growth observed for each strain. All liquid cultures were grown at 25 °C and shaken at 165 rpm. Cells were harvested by centrifugation at 8,000 x g for 10 min at 4 °C, washed with five mL of 0.5× Murashige & Skoog media (MS), and resuspended in 12 mL of 0.5× MS. Since insufficient biomass could be obtained from all strains to hit the target final OD_600_ of 0.01, the inoculum was produced by diluting all strains into 0.5× MS to a final OD_600_ between 0.0005-0.01 depending on species biomass production (Table 1). A heat inactivated consortium (KC) inoculum was prepared by boiling the inoculum at 100 °C for 1.5 h. This treatment was intended to eliminate bacterial viability while retaining bacterial biomass as a control for the live consortium. Samples of both live and inactivated inoculum were saved at −20 °C for amplicon sequencing.

**Table 1.** Growth schedule of the selected strains for the preparation of consortium inoculum. Classification based on the Type (Strain) Genome Server showing digital DNA-DNA hybridization (dDDH) Genome BLAST Distance phylogeny formula 2 (d4). Isolates were inoculated on R2A plates for 2-4 days (Days on R2A), then transferred to tryptic soy broth (TSB) at the indicated strength. Growth time of starter culture (five mL) and final culture (30 mL). Final optical density at 600 nm of each strain in the inoculum (Final OD).

| Colony ID | Isolate ID | Closest typed species | dDDH (d4, in %) | Days on R2A | TSB strength | Hrs in starter culture | Hrs in final culture | Final OD |
|---|---|---|---|---|---|---|---|---|
| CT11-1 | Prar1 | Priestia aryabhattai JCM 13839 | 73.6 | 2 | 1x | 14 | 16 | 0.01 |
| CT11-16 | Pseusp16 | Pseudomonas jessenii DSM 17150 | 46.8 | 2 | 1x | 14 | 16 | 0.0095 |
| CT11-20 | Pseusp20 | Arthrobacter liuii CGMCC1.12778 | 32.1 | 3 | 0.1x | 14 | 16 | 0.0062 |
| CT11-24 | Chor24 | Chryseobacterium oranimense DSM 19055 | 81.7 | 3 | 1x | 14 | 16 | 0.008 |
| CT11-25 | Dugasp25 | Rugamonas aceris SAP-35 | 35.9 | 3 | 0.1x | 16 | 30 | 0.0076 |
| CT11-34 | Lutesp34 | Luteibacter jiangsuensis CGMCC 1.10133 | 46.1 | 4 | 0.1x | 16 | 40 | 0.0105 |
| CT11-41 | Varisp41 | Variovorax boronicumulans NBRC 103145 | 87.7 | 4 | 0.1x | 14 | 40 | 0.012 |
| CT11-44 | Dugasp44 | Duganella vulcania FT81W | 61.3 | 3 | 0.1x | 16 | 30 | 0.0038 |
| CT11-53 | Varisp53 | Variovorax gossypii DSM 100435 | 37.8 | 4 | 1x | 16 | 40 | 0.0075 |
| CT11-57 | Curtsp57 | Curtobacterium pusillum ATCC 19096 | 55.6 | 4 | 1x | 14 | 14 | 0.003 |
| CT11-59 | Rugasp59 | Rugamonas rivuli FT103W | 87.8 | 3 | 1x | 16 | 72 | 0.007 |
| CT11-66 | Sphisp66 | Sphingomonas kyeonggiensis DSM 101806 | 41.2 | 4 | 0.1x | 24 | 30 | 0.0024 |
| CT11-68 | Paph68 | Paraburkholderia phenoliruptrix LMG 22037 | 78.6 | 3 | 1x | 16 | 30 | 0.0009 |
| CT11-81 | Flavsp81 | Flavobacterium pectinovorum DSM 6368 | 26.3 | 4 | 0.1x | 48 | 48 | 0.0013 |
| CT11-82 | Rhizsp82 | Rhizobium croatiense 13T | 37.1 | 4 | 0.1x | 30 | 48 | 0.0043 |
| CT11-83 | Polasp83 | Polaromonas jejuensis NBRC 106434 | 24.5 | 4 | 0.1x | 30 | 30 | 0.0014 |
| CT11-86 | Mucisp86 | Mucilaginibacter rubeus CGMCC 1.15913 | 41.6 | 4 | 0.1x | 30 | 30 | 0.0005 |
| CT11-90 | Panico90 | Paenarthrobacter nicotinovorans DSM 420 | 88.8 | 3 | 1x | 14 | 14 | 0.0101 |
| CT11-92 | Mace92 | Massilia cellulosiltytica NEAU-DD11 | 83.4 | 4 | 0.1x | 16 | 16 | 0.0023 |
| CT11-105 | Koco105 | Kosakonia cowanii JCM 10956 | 72.1 | 3 | 0.1x | 14 | 14 | 0.0091 |
| CT11-110 | Ralssp110 | Ralstonia wenshanensis 56D2T | 63.5 | 3 | 0.1x | 16 | 30 | 0.005 |
| CT11-111 | Paensp111 | Paenibacillus zeisoli 3-5-3 | 34.8 | 4 | 1x | 30 | 14 | 0.0044 |
| CT11-113 | Stgeni113 | Stenotrophomonas geniculata JCM 13324 | 81 | 3 | 1x | 14 | 14 | 0.0101 |
| CT11-119 | Hefr119 | Herbaspirillum frisingense GSF30 | 82.5 | 3 | 0.1x | 30 | 30 | 0.0016 |
| CT11-120 | Padi120 | Pantoea dispersa CCUG 25232 | 83.9 | 3 | 1x | 14 | 14 | 0.0099 |
| CT11-132 | Azosp132 | Azospirillum oryzae COC8T | 54.8 | 3 | 1x | 30 | 30 | 0.0005 |
| CT11-134 | Mesosp134 | Mesorhizobium carmichaelinearum ICMP 18942 | 39.9 | 4 | 0.1x | 48 | 72 | 0.0012 |

Seeds were surface sterilized as described above and grown gnotobiotically in 42 oz (1.24 L) Whirl-Pak® stand-up bags (Nasco Sampling, Pleasant Prairie, USA) containing 150 mL of sterile calcined clay (Pro’s choice rapid dry, Oil-Dri Corporation of America, Chicago, USA) and 60 mL of 0.5× MS media containing the corresponding consortium inoculum (Wagner *et al*., 2021). The live consortium (LC) was inoculated into four wheat varieties: Catawba (n=6), Hilliard (n=6), Shirley (n=5), and USG3640 (n=6); each genotype was also inoculated with a no consortium control (NC, n=6 per genotype). The KC treatment was only applied to cultivar Catawba (n=6). The bags were sealed with AeraSeal^TM^ sealing film (Excel Scientific, Victorville, USA). Plants were grown up to 15 days post emergence (DPE) with a 12 h photoperiod and day/night temperatures of 22/16 °C. Destructive sampling of Catawba occurred at 5, 10, and 15 DPE. Destructive sampling only occurred on DPE 15 for the other varieties. The effective quantum yield of photosystem II was measured as an indicator of chlorophyll fluorescence using a MultispeQ v1.0 (PhotosynQ, USA) under ambient light conditions. Due to small size of the plants, two leaves were positioned together within the leaf clamp to cover the measurement area. Measurements were performed using the Photosynthesis Rides 2.0 protocol and processed in PhotosynQ Software V1.10.59. Roots and shoots were separated using sterile scissors. Roots were harvested and cleaned as described above and blot dried. Shoots and clean roots were weighed to determine fresh biomass. Clean roots were cut into 0.5 cm pieces and stored at −20 °C until further processing.

### DNA extraction and amplicon sequencing of inoculated wheat roots

DNA was extracted from clean wheat roots inoculated with consortium (live and heat-inactivated) and no consortium and initial root slurry with the DNeasy Plant Pro Kit (Qiagen, Hilden, Germany). The 16S rRNA gene variable regions V3-V4 were amplified using primer pair 341F-860R (Hansen *et al*., 2012) by the two-step PCR described by Thøgersen *et al*. (2024). Libraries were equimolarly pooled and sequenced on an Illumina MiSeq using the MiSeq Reagent Kit V2 500-cycle (Illumina, San Diego, USA) for 2 × 250 bp paired-end (PE) reads. Sequencing yielded a total of 7,544,224 reads with a per sample average of 81,121 reads. Data were demultiplexed and adapter were removed from paired-end reads using Cutadapt in QIIME 2 v. 2022.8 (Bolyen *et al*., 2019) allowing adapter matching anywhere within reads (forward: TCGTCGGCAGCGTCAGATGTGTATAAGAGACAG; reverse: GTCTCGTGGGCTCGGAGATGTGTATAAGAGACAG). On average 37,150 PE reads were obtained per sample. In samples with no consortium and KC approximately 99 ± 0.81 % of the reads matched plant mitochondrial and chloroplast sequences. Due to the small amounts of bacterial reads in these samples, they were removed from the amplicon analysis as they were considered to have negligible bacterial colonization. In contrast, in samples with consortium, on average 27 ± 22 % of the reads matched the plant sequences. Two different strategies were used to analyze data from the initial root slurry and samples from the consortium-inoculated plants. Reads from the initial root slurry were filtered, denoised, curated, and chimera checked with DADA2 v 1.16 (Callahan *et al*., 2016) and LULU v 0.1.0 (Frøslev *et al*., 2017). Taxonomy was assigned using the SILVA v138 database (silva_nr99_v138.1_wSpecies_train_set). For the remaining samples, reads were quality filtered and merged using bbsplit v 38.07 (Bushnell, 2014). Merged reads were then mapped to a custom-made database using a minimum sequence identity of 99% to obtain the community composition of consortium inoculated wheat plants. The custom-made database contained the 16S rRNA gene sequences of all consortium members and was supplemented with the 16S rRNA gene sequences from wheat mitochondria, chloroplast, and plastids within the SILVA database. These wheat sequences were clustered at 99% similarity using cd-hit-est (Fu *et al*., 2012) prior to analysis. Amplicon sequencing data is available from NCBI archives BioProject PRJNA1415243.

### SynCom member candidate selection criteria

Selection criteria were defined to guide strain inclusion based on measurable features derived from amplicon sequencing data and knowledge of plant-associated microbiome dynamics. Because mutants of colonization related genes reduces colonization efficiency and associated beneficial traits (Chin-A-Woeng *et al*., 2000; Cole *et al*., 2017), robust colonizers were identified as strains that increased in relative abundance in wheat roots compared to the original consortium inoculum and exceeded a minimum relative abundance threshold of 0.01%. Because differentially abundant taxa have been associated with plant benefits (Kwak *et al*., 2018; Lazcano *et al*., 2021), strains potentially responsive to host or environmental cues were further evaluated based on changes in relative abundance across time points and between cultivars. Because core taxa may reflect conserved functions regarding host associations and keystone taxa may play important roles in maintaining community structure and function (Toju *et al*., 2018; Risely, 2020), ecologically relevant taxa were identified as members of the shared or temporal core, as well as taxa previously reported as core of keystone members of the wheat microbiome. Based on these considerations, candidate strains were evaluated according to five criteria: 1) increased relative abundance in roots compared to the inoculum, 2) relative abundance of 0.01% in root samples, 3) variation in abundance across time points or cultivars, 4) classification as temporal or cultivar core members (at ≥90% prevalence), and 5) prior identification as a core or keystone taxon in the literature. Strains meeting at least three of these criteria were considered for inclusion in the SynCom.

### Statistical analysis

All statistical analyses were performed using R statistical software version 4.3.0 (R Core Team 2023). One-sample t-tests were performed to compare the relative abundances of each SynCom member at 15 DPE to their respective relative abundances in the initial inoculum as a fixed reference; *p-values* were adjusted using the Benjamini-Hochberg (BH) correction for multiple testing at α = 0.1. The effect of the consortium on wheat phenotype was evaluated at 15 DPE in a two-way analysis of variance (ANOVA) with treatment, genotype, and their interaction as fixed effects at a level of *p* <0.05. Model residual diagnostics were evaluated using residual plots, the Shapiro-Wilk test for normality, and Levene’s test for heteroscedasticity. All analyses were performed on raw data with the exception of root-to-shoot ratio, which was log-transformed to normalize residual distribution. Multiple comparisons were made using estimated marginal means using the emmeans (Lenth & Piaskowski, 2025) package with Tukey *p-value* adjustment. Differences in community composition were assessed using Euclidean distances on centered log-ratio (CLR) transformed abundances following a pseudocount addition (x + 0.02), using the following R packages: phyloseq (McMurdie & Holmes, 2013), compositions (Boogaart *et al*., 2025), and vegan (Oksanen *et al*., 2025). The effect of genotype or harvest day on community composition were evaluated using the randomized residual permutation procedure (RRPP) using the lmrrpp function (Collyer & Adams, 2018), using Type III sums of squares. Homogeneity of multivariate dispersion was evaluated using betadisper function. Differentially abundant strains across wheat genotypes and harvest days were examined using the DESeq2 package (Love *et al*., 2014) with BH adjustments for multiple comparisons (α=0.1). Data visualization was performed using the ggplot2 (Wickham, 2016) package and the graphical abstract was created in Biorender.com. Assistance with code development and data curation for statistical analyses was provided by ChatGPT 5.6 (OpenAI, San Francisco, CA), which was used to streamline table formatting and prepare scripts for data processing.

## Results

### Isolation strategies resulted in the recovery of taxa identified as part of the wheat core microbiota

To recover a diverse set of wheat-associated bacteria, we employed two isolation strategies with four cultivation media each (VXylG, R2A 1/10, 1/20, 1/100). Isolates were plated after either one (direct plating strategy) or two (host-mediated plating strategy) rounds of host selection in which the plant acted as a selective filter. The isolation effort initially yielded 125 pure isolates encompassing 31 genera and 53 species (Table S2). Seventy-three isolates were obtained by direct plating, and 52 isolates were obtained using the host-mediated approach. *Priestia, Peribacillus*, *Gottfriedia, Chryseobacterium, Arthrobacter, Flavobacterium, Mucilaginibacter, Variovorax, Agrobacterium, Luteibacter, Paraburkholderia, Polaromonas,* and *Roseateles (Mitusaria)* were obtained only by direct plating, whereas *Burkholderia, Stenotrophomonas, Telluria (Massilia), Pantoea, Achromobacter*, *Mesorhizobium, Azospirillum, Herbaspirillum, Kosakonia,* and *Ralstonia,* were isolated only in the host-mediated strategy. This could indicate that the combination of both strategies, direct and host-mediated, leads to greater diversity recovery.

To assess the coverage of our isolation efforts, we determined the community composition of the original root slurry through 16S rRNA gene amplicon sequencing and compared it to the full length 16S rRNA gene sequences of isolates. We obtained 90 amplicon sequence variants (ASVs) in the original root slurry after removal of plant-associated and lowly abundant sequences (< 0.01%). ASVs were assigned to 15 phyla, of which Pseudomonadota, Actinomycetota, and Bacteroidota were the most abundant, encompassing 89% of the sequencing reads (Figure 1a) with read abundances between 18.5% and 46.8%. Myxococcota, Fibrocacterota, Chloroflexota, and Bacillota constituted an additional 7.5% of the sequencing reads in the community with each phylum’s abundance between 3.4% and 1.1%. The remaining eight phyla had read abundances below 1%, accounting for the remaining 3.5% of the community. Consistent with the structure of this root slurry community, isolate sequences were classified into four phyla, with Pseudomonadota and Actinomycetota encompassing 87.9% of the collection (Figure 1b); isolates within Bacillota and Bacteroidota made up the remaining 12.1% of the collection.

**Figure 1.**
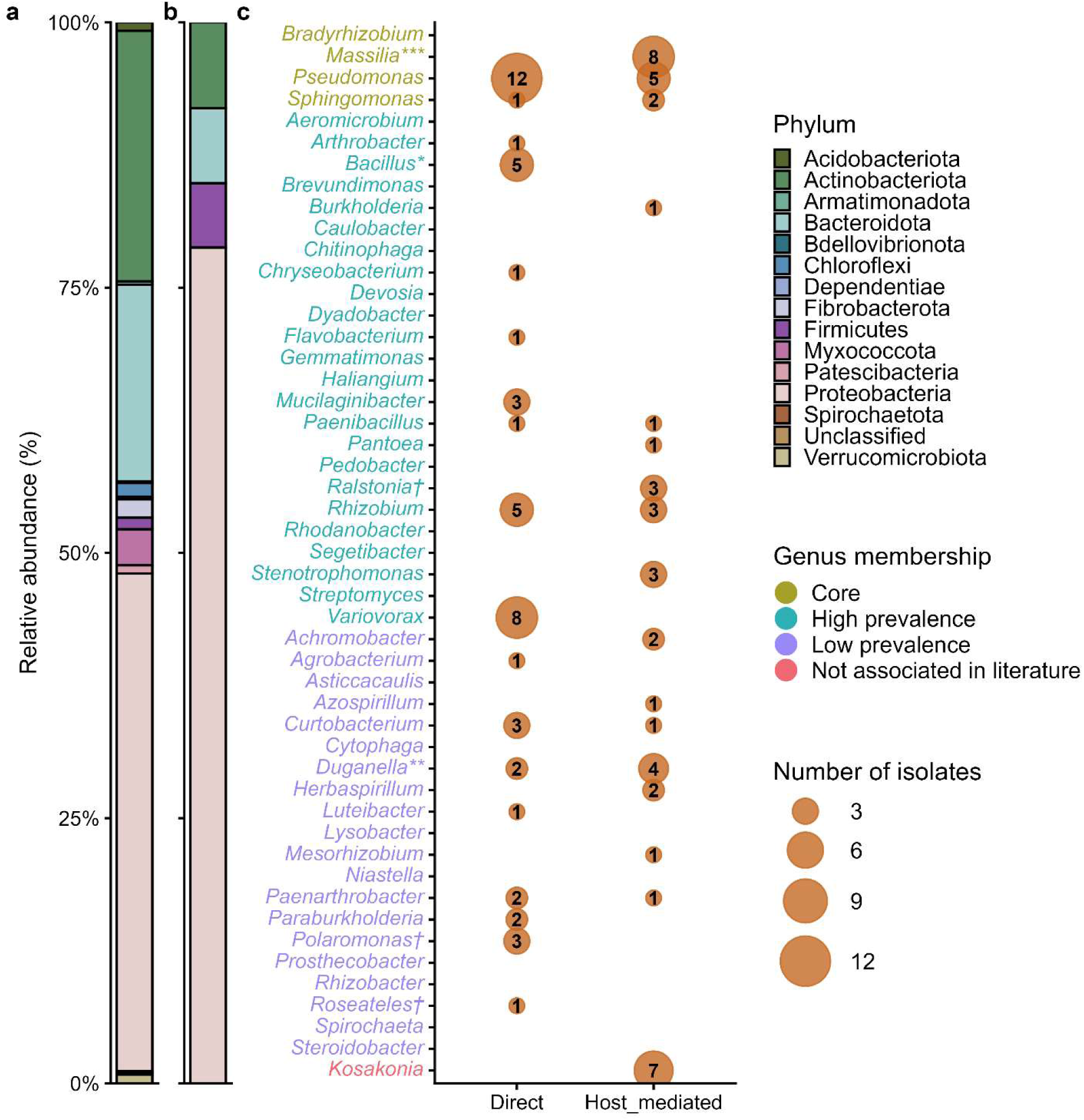
Taxonomic composition of the wheat root microbiome and its cultured isolates. (**a**) Relative abundances of bacterial phyla in the original root slurry, determined by amplicon sequencing of the 16S rRNA gene. Values represent relative abundances (%) calculated from sequencing reads. (**b**) Phylum-level composition of recovered isolates, based on 16S rRNA gene sequences from individual isolates. Relative abundances (%) reflect the proportion of isolates assigned to each phylum. (**c**) Recovery of genera identified as wheat-associated in Kavamura *et al*. (2021) classified as core when shared in at least 60% of studies included (green), highly prevalent as taxa shared in at least 40 % of studies (turquoise), low prevalence taxa as those shared in less than 40% of studies (purple), or not mentioned in reference list or literature as taxa not associated in literature (pink). Bubble size indicates the number of isolates recovered per genus in each isolation strategy (direct plating or host-mediated enrichment). Closely related genera potentially undifferentiated via amplicon sequencing grouped together: * includes *Priestia, Peribacillus*, and *Gottfrieda*, **includes *Rugamonas*, and ***includes *Telluria*. †Taxa not found in the reference list, but present in other wheat microbiome studies.

At the genus level, ASVs in the root slurry were assigned to 74 distinct taxonomic groups, out of which only 47 could be classified at the genus level (Table S3). The most abundant genera in the root slurry were *Streptomyces, Polaromonas, Chitinophaga, Niastella, Sphingomonas*, and *Kineosporia*; these taxa encompassed 50% of the sequencing reads with abundances between 5 and 16.4%. Of these 47 genera in the root slurry, 27.5% were recovered by our isolation effort, including the highly abundant *Polaromonas* and *Sphingomonas.* Also isolated were members of the genera *Mesorhizobium, Duganella, Allorhizobium-Neorhizobium-Pararhizobium-Rhizobium*, and *Flavobacterium*, which accounted for 9.2 % of the original root slurry sequencing reads. The isolation effort also recovered four genera that were lowly abundant in the root slurry (2.8% of reads), *Mucilaginibacter, Pseudomonas, Bacillus,* and *Burkholderia-Caballeronia-Paraburkholderia*, as well as sixteen genera that were not detected by amplicon sequencing: *Variovorax, Massilia, Kosakonia, Curtobacterium, Ralstonia, Chryseobacterium, Paenarthrobacter, Stenotrophomonas, Achromobacter, Herbaspirillum, Paenibacillus, Pseudarthrobacter, Luteibacter, Mitsuaria, Pantoea*, and *Azospirillum*.

To assess the taxonomic coverage of our collection relative to previously reported wheat-associated taxa, we leveraged the list of 256 wheat-associated genera compiled by Kavamura *et al*. (2021) from 10 studies encompassing eight countries across four continents. From this list we considered 27 of these genera highly prevalent (Table S4). Our isolates covered 51.8% of the highly prevalent genera, including core taxa such as *Sphingomonas, Pseudomonas,* and *Massilia* (Figure 1c). Moreover, the collection includes another 10 genera associated with the wheat microbiome but at lower prevalence (less than 40% of studies). In addition, we obtained isolates from *Kosakonia, Polaromonas, Ralstonia, and Roseateles (Mitsuaria),* which are not within the 256 previously mentioned wheat-associated taxa. However, *Ralstonia, Roseateles (Mitsuaria)* and *Polaromonas* have been associated with wheat in other studies (Yin *et al*., 2013; Ofek-Lalzar *et al*., 2014; Rascovan *et al*., 2016; Gdanetz & Trail, 2017; Pagé *et al*., 2019; Bakker & McCormick, 2019; Johnston-Monje *et al*., 2021; Garrido-Sanz & Keel, 2025), highlighting the non-exhaustive nature of the reference list.

### Broad presence of genes associated with plant–microbe interactions within the isolate collection

To catalog important plant–microbe interaction genes found within isolates, we performed whole genome sequencing of 100 isolates, selected out of the 125 initial isolates (Fig S1). Short read sequencing resulted in genome assemblies with contig numbers ranging from 16 to 2,759 with N50 values between 2,478 and 698,027. The genomes had an average completeness of 99.1%, and the contamination for all but one genome assembly was below 10%. The highly contaminated genome was therefore removed from the collection and further analyses resulting in 99 high-quality genomes. Representative genomes selected for subsequent long read sequencing and hybrid genome assembly, resulted in 63 complete genomes. The final assemblies, including short-read genome assemblies, had total lengths between 3.7 to 8.6 Mbp, and the GC content ranged between 33 and 71% (Table S5). Contig number ranged between 1 and 2,038 with N50 values between 4,375 and 7,318,753. Final assemblies had an average genome completeness of 99.8% with a contamination average of 0.6%. The average proportion of Benchmarking Universal Single-Copy Orthologs (BUSCOs) found was 92% and the average gene density was 0.9. The 99 genomes were closely related to 53 type strains and classified into 31 genera (Table S6). Forty-eight isolates were identified as potential new species after removal of redundant genomes. We have made 88 strains, representative of all 31 genera recovered in this isolation effort, publicly available at the DSMZ.

To facilitate the use of this bacterial collection by the wider scientific community, we screened all genomes against a literature derived target list of genes associated with plant-microbial interactions to provide an overview of the functional potential of this resource. The target list contained 4,843 KOs categorized into 27 broad categorical functions and 93 additional specific functions. These features were widely distributed across the collection, with an average of 1487 KOs identified per genome (Table S7). Strains within *Pseudomonas, Burkholderia* and *Paraburkholderia* exhibited the highest gene counts per genome (average: 1737), whereas *Curtobacterium, Flavobacterium*, and *Chryseobacterium* showed the lowest (average: 899). The distribution of KOs across the collection broadly reflects phylogenetic relationships, while differences in predicted functional repertoire among taxa suggest varying degrees of functional breadth, potentially reflecting generalist and specialist strategies. To facilitate visualization of these patterns, a subset of 42 functions was selected to represent key processes involved in plant growth promotion, nutrient cycling, and adaptations to the plant environment and summarized their distribution by calculating the proportion of genes present within each genome for each functional category (Figure 2). While many genes linked to plant colonization traits, plant growth promotion, and nutrient cycling were widespread, some more specialized functions were restricted to specific phylogenetic groups or strains. For example, N fixation genes were only present in three isolates within the collection, Pseusp138 (*Pseudomonas* sp.), Azosp132 (*Azospirillum* sp.) and Paensp38 (*Paenibacillus* sp.). Additionally, we assessed the biosynthetic potential of the collection by identifying biosynthetic gene clusters (BGCs). A total of 568 BGCs were identified in the collection, representing 60 cluster classes (Table S8). The most abundant classes identified included terpenes, unspecified ribosomally synthesized and post-translationally modified peptide products, aryl polyenes, non-ribosomal peptide synthases, redox cofactors, siderophores, betalactones, type III polyketide synthase, and homoserine lactones (Fig S2). These BGCs highlight the collection’s potential to produce diverse secondar metabolites, including compounds associated with antimicrobial and cytotoxic compounds relevant to pathogen control in plants (Table S9). Distribution patterns of functional traits and secondary metabolite production are discussed in more detail in the Supplementary Information.

**Figure 2.**
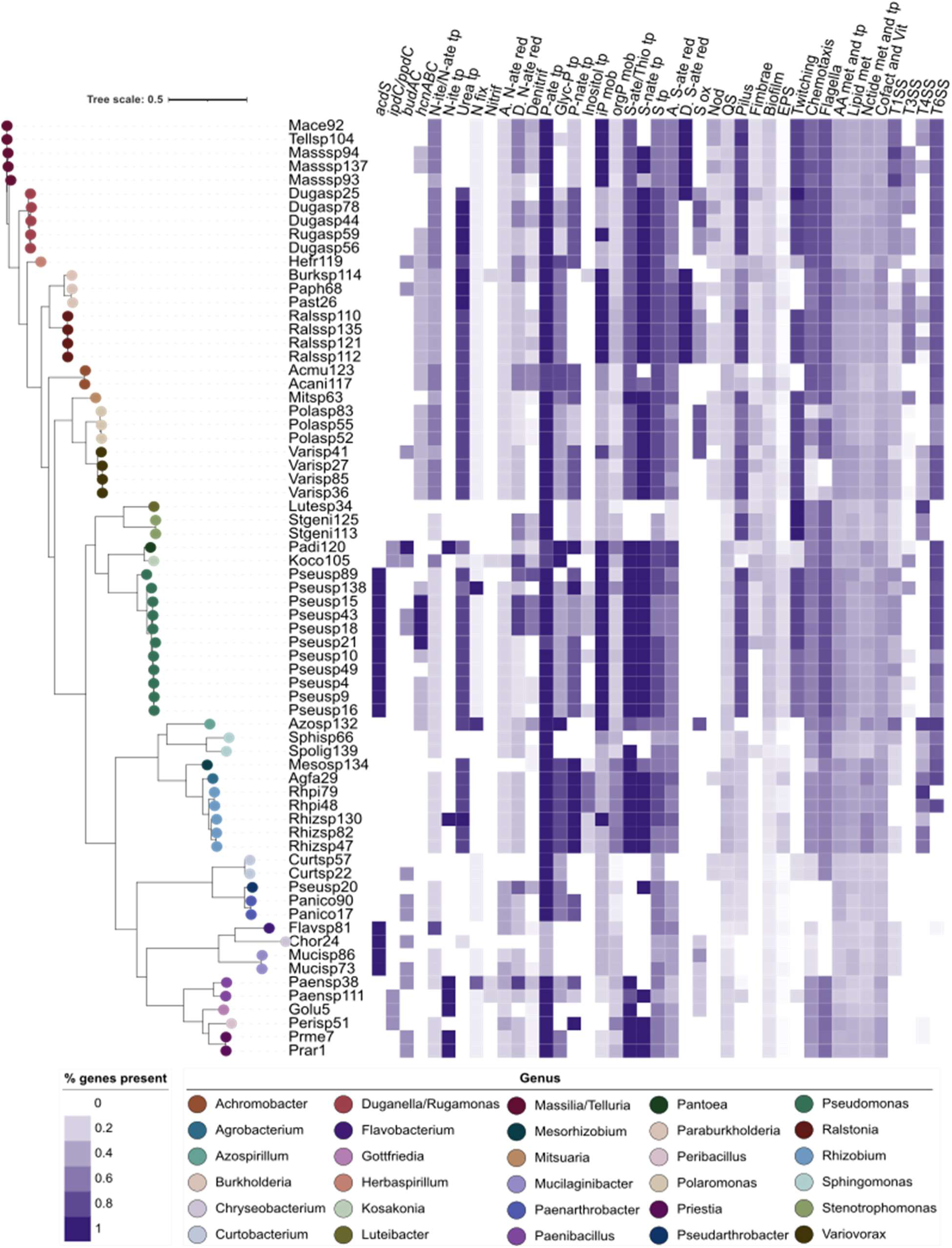
Functional gene distribution across 69 representative genomes from the TriMic collection available at the DSMZ. Representative genomes were selected from 88 isolates based on phylogenetic relationships, prioritizing genomes completeness and hybrid assemblies when available Functional profiles for all 88 isolates are provided in Supplementary Table 7. The heatmap depicts the proportion of target KEGG orthologs detected per genome within each of the selected functional categories, calculated as the number of genes present in the genome relative to the total number of genes in the category (n= number of genes in category). Functional categories are as follows: Plant growth promotion: 1-aminocyclopropane-1-carboxylate deaminase (*acdS*, n = 2), indole-3-acetic acid synthesis (*ipdC/ppdC*, n = 2), acetoin/2,3-butanediol synthesis (*budAC*, n = 2), and hydrogen cyanide production (*hcnABC*, n = 5). Nitrogen and phosphorus cycling: nitrate/nitrite transport (N-ite/N-ate tp, n = 5), nitrite transport (N-ite tp, n = 1), urea transport (Urea tp, n = 6), nitrogen fixation (N fix, n = 11), nitrification (Nitrif, n = 5), assimilatory nitrate reduction (A. N-ate red, n = 5), dissimilatory nitrate reduction (D. N-ate red, n = 7), denitrification (Denitrif, n = 13), phosphate transport (P-ate tp, n = 6), glycerol-phosphate transport (Glyc-P tp, n=5), phosphonate transport (P-nate tp, n = 4), inositol transport (Inositol tp, n = 9), inorganic phosphorus mobilization (iP mob, n = 6), and organic phosphorus mobilization (orgP mob, n = 21). Adaptation to plants and colonization: sulfate/thiosulfate transport (S-ate/Thio tp, n=4), sulfonate transport (S-nate tp, n = 3), sulfur relay system (SRS, n = 5), assimilatory sulfate reduction (A. S-ate red, n = 7), dissimilatory sulfate reduction (D. S-ate red, n = 1), sulfur oxidation (S ox, n = 4), nodulation (Nod, n = 4), quorum sensing (QS, n = 68), pilus formation (Pilus, n = 8), fimbriae (Fimbriae, n = 6), biofilm formation (Biofilm, n = 85), exopolysaccharide biosynthesis (EPS, n = 36), twitching motility (Twitching, n=6), chemotaxis (Chemotaxis, n = 27), flagellar motility (Flagella, n = 42), amino acid metabolism and transport (AA met and tp, n=451), lipid metabolism (Lipid met, n = 107), nucleotide metabolism and transport (Nctide met and tp), metabolism of cofactors and vitamins (Cofact and Vit), and secretion systems including Type I (T1SS, n = 6), Type III (T3SS, n = 24), Type IV (T4SS, n = 19), and Type VI (T6SS, n = 20).

### Inoculation with 27 strains revealed temporal shifts in community structure

To guide construction of a reduced SynCom, we evaluated the effect of a 27-member consortium spanning the phylogenetic diversity of the recovered isolates on plant performance and tracked its community composition across four wheat cultivars over two weeks using amplicon sequencing. There was no significant effect of the consortium on any of the plant performance metrics measured, and the cultivar was the main driver of plant phenotype differences (Figure S3, Table S10), thus indicating consortium compatibility with the host and the absence of overt pathogenic effects.

To assess which isolates were robust root colonizers, we compared the change in abundance of each species in wheat roots after two weeks of growth to the initial inoculum. When compared to the initial inoculum, 21 strains had significantly lower relative read abundances within plant roots (Figure 3). Only four strains had significantly greater relative read abundances when compared to the inoculum (Table S11). Strains Paph68, Masssp92, Curtsp57, and Ralsp110 increased significantly in abundance by 7000%, 480 %, 227%, and 367%, respectively whereas Flavsp81, Varisp53, Prar1 became undetectable. Differences in the community composition through time were assessed at three different harvest days 5, 10, and 15 days post emergence (DPE) in cultivar Catawba (Figure 3a) and differences between cultivars were compared at 15 DPE only (Figure 3b). Harvest day had a significant effect on the community composition (RRPP F_2,14_= 1.97, R^2^ = 0.219, *p-value* = 0.015). There were significant differences in Catawba root community composition between 5 DPE and 10 DPE (*p-value* = 0.028). Dispersion analysis (*p-value* =0.904) confirmed that this effect is a result of difference in community centroids and not in dispersion. We determined the differentially abundant (DA) strains between time points. There were four species that changed significantly over time (Table S12). Overall, Lutesp34, and Stgeni113 decreased over time, whereas Dugasp44 increased. Koco105, on the other hand, showed a decrease from day 5 to 10 but increased again on day 15 to read abundances equivalent to those at day 5 (Fig S4).

**Figure 3.**
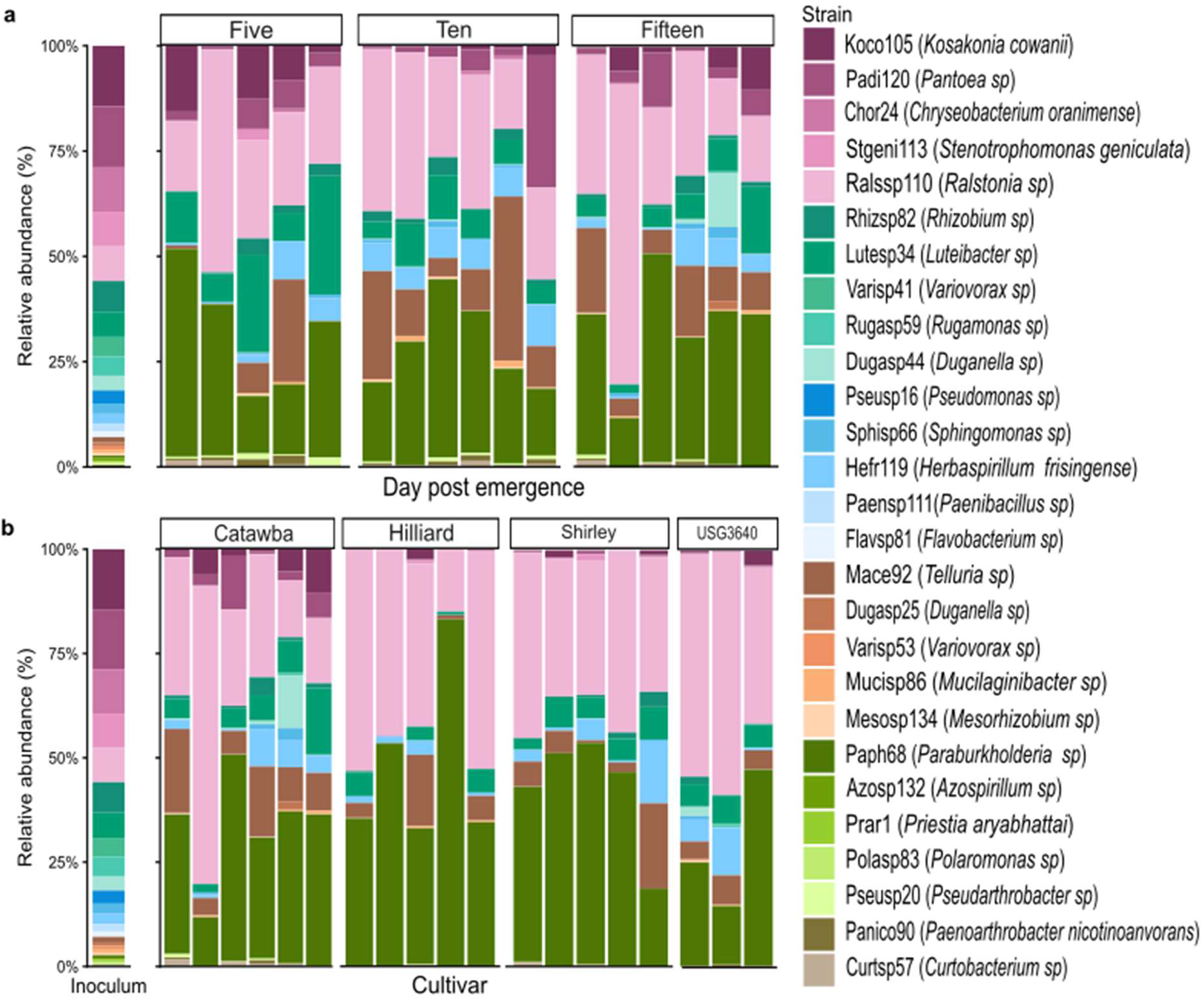
Microbial community composition dynamics on roots from four wheat cultivars inoculated with a 27-member consortium. Relative abundances of consortium members are expressed as percentages based on 16S rRNA gene amplicon sequencing reads in individual samples. The inoculum composition is shown as a reference: **a.** Temporal dynamics of community composition in cultivar Catawba at five (n=5), ten (n=6), and fifteen (n=6) days post emergence, **b.** Cultivar-specific community composition at fifteen days post emergence across four wheat cultivars, Catawba (n=6), Hilliard (n=5), Shirley (n=5), USG3640 (n=3).

Cultivar had no effect on the overall community composition (RRPP, F_3,15_= 1.219, R^2^= 0.196, *p-value* = 0.171,). However, we observed that ten strains had significant differential abundances between cultivars (Table S13). The most differentially abundant strains were found between Catawba and the other three cultivars; where Mucisp86, Rhisp82, Koco105, Padi120, Dugasp25, Dugasp44, Pseusp16, Sphisp66, Varisp41 and Curtsp57 had greater abundances in Catawba in comparison to other cultivars (Fig S5). Dugasp44 was also differentially abundant between cultivars USG3640 and Hilliard, where its abundance was greater in USG3640 than Hilliard. Between Shirley and Hilliard, Mucisp86 and Rhizsp82 had greater and lower abundance, respectively, in Shirley than in Hilliard. We also determined the shared core across time and cultivar at a prevalence rate of 90% with a minimum abundance of 0.01%. Eleven species were identified as the temporal core; Ralssp110, Mace92, Hefr119, Rhizsp82, Koco105, Pseusp20, Padi120, Panico90, Paph68, Curtsp57 and Lutesp34. In contrast, only six strains were identified as core between wheat cultivars at the same prevalence rate, Ralssp110, Mace92, Hefr119, Padi120, Paph68, and Lutesp34.

### The proposed wheat root SynCom captures wheat core taxa and preserves functional diversity of the collection

The inclusion of strains in the final SynCom was determined based on five criteria (Table 2) designed to capture four key attributes: root colonization ability, dynamic behavior indicative of activity, taxonomic representation and ecological relevance within the wheat microbiome. Fifteen strains met at least three criteria and were considered for the SynCom. These strains span 14 genera, with one strain per genus except for *Duganella*, which yielded two candidates. To reduce potential intra-species competition, Dugasp25 was selected as a single candidate from this genus due to its potential to produce siderophores and homoserine lactones. Even though Azosp132 did not meet the minimum candidate criteria; this strain was included due to its potential for nitrogen fixation to complement the functional capabilities of the SynCom as a whole. The proposed SynCom consists of 15 species from 15 genera: Koco105, Ralssp110, Hefr119, Padi120, Azosp132, Pseusp16, Dugasp25, Lutesp34, Varisp41, Curtsp57, Sphisp66, Paph68, Rhizsp82, Mucisp86, and Mace92 (Table 2). This SynCom also captures approximately 55% of the most prevalent genera reported across wheat microbiome studies and includes taxa previously identified as core and keystone members. In addition to taxonomic representation, the SynCom retains the functional breadth of the isolate collection, including diverse plant-microbe interaction functions and secondary metabolite BGCs (Fig S6 5).

**Table 2.** Criteria used for selection of strains included in the wheat root synthetic community (SynCom). Strains in the 27-member consortium were considered for SynCom candidacy if three out of the five criteria were met. Selection criteria were based on: (1) percent relative read abundances in wheat roots above 0.01 %, (2) increase of relative abundance compared to inoculum; (3) differential abundance across time points or between cultivars, as determined using DESeq2; (4) shared core membership across harvest points or cultivars at 90% prevalence of samples and (5) prior identification as core or keystone taxa in the literature. Isolate IDs in bold identify strains meeting the minimum threshold for SynCom candidacy. * Denotes inclusion, despite meeting only 2/5 criteria to supplement functional capabilities of the SynCom.

| Isolate ID | Genus | > 0.01 % abundance | Increase from inoculum | Differentially abundant | Prevalence core at 90% | Core/Keystone in literature |
|---|---|---|---|---|---|---|
| Prar1 | *Priestia* |  |  |  |  |  |
| **Koco105** | *Kosakonia* | Yes |  | Yes | Yes |  |
| **Ralssp110** | *Ralstonia* | Yes | Yes |  | Yes |  |
| Paensp111 | *Paenibacillus* |  |  |  |  | Yes |
| Stgeni113 | *Stenotrophomonas* | Yes |  | Yes |  |  |
| **Hefr119** | *Herbaspirillum* | Yes |  |  | Yes | Yes |
| **Padi120** | *Pantoea* | Yes |  | Yes | Yes | Yes |
| Azosp132* | *Azospirillum* | Yes |  |  |  | Yes |
| Mesosp134 | *Mesorhizobium* | Yes |  |  |  |  |
| **Pseusp16** | *Pseudomonas* | Yes |  | Yes |  | Yes |
| Pseusp20 | *Arthrobacter* | Yes |  |  | Yes |  |
| Chor24 | *Chryseobacterium* | Yes |  |  |  | Yes |
| **Dugasp25** | *Duganella* | Yes |  | Yes |  | Yes |
| **Lutesp34** | *Luteibacter* | Yes |  | Yes | Yes | Yes |
| **Varisp41** | *Variovorax* | Yes |  | Yes |  | Yes |
| **Dugasp44** | *Duganella* | Yes |  | Yes |  | Yes |
| Varisp53 | *Variovorax* |  |  |  |  | Yes |
| **Curtsp57** | *Curtobacterium* | Yes | Yes | Yes | Yes |  |
| Rugasp59 | *Rugamonas* | Yes |  |  |  | Yes |
| **Sphisp66** | *Sphingomonas* | Yes |  | Yes |  | Yes |
| **Paph68** | *Paraburkholderia* | Yes | Yes |  | Yes |  |
| Flavsp81 | *Flavobacterium* |  |  |  |  | Yes |
| **Rhizsp82** | *Rhizobium* | Yes |  | Yes | Yes |  |
| Polasp83 | *Polaromonas* |  |  |  |  |  |
| **Mucisp86** | *Mucilaginibacter* | Yes |  | Yes |  | Yes |
| Panico90 | *Paenoathrobacter* | Yes |  |  | Yes |  |
| **Mace92** | *Telluria (Massilia)* | Yes | Yes |  | Yes | Yes |

## Discussion

Here, we describe the development of a resource for dissecting wheat-microbe interactions consisting of a publicly available, curated culture collection of wheat root-associated bacteria and a proposed reduced complexity SynCom assembled from strains in the collection. Key to the utility of these resources is how well they capture the taxonomic and functional diversity of the host microbiota(Northen *et al*., 2024). The TriMic collection is composed of taxa generally associated with the wheat microbiota including highly prevalent and core taxa. The isolates in this collection belong to four main phyla, Pseudomonadota, Bacteroidota, Actinomycetota, and Bacillota, which constitute roughly 50% of the wheat microbiome composition in culture independent studies (Gdanetz & Trail, 2017; Mavrodi *et al*., 2018; Araujo *et al*., 2019; Chen *et al*., 2019; Schlatter *et al*., 2020; Rossmann *et al*., 2020; Simonin *et al*., 2020; Kavamura *et al*., 2021). This taxonomic distribution at the phylum level is not only consistent with other isolate collections from maize, wheat, rice, and Medicago (Dai *et al*., 2025; Garrell *et al*., 2026b), but also consistent with the known plant-microbiome (Trivedi *et al*., 2020), highlighting the relevance of this collection to plant-microbiome studies.

Focusing on highly prevalent and core microbiota provides a practical framework to assess taxonomic coverage of microbial collections (Busby *et al*., 2017), because culture-dependent and culture-independent methods often lead to discrepancies in community composition (Kavamura *et al*., 2019; Becker *et al*., 2024) and methodological biases and environmental variability confound genus level comparability between microbiome studies (Dastogeer *et al*., 2020; Forry *et al*., 2024). Core microbiota comprises microbes consistently associated with a host species across habitats and that might reflect critical relationships that can be leveraged for enhanced plant productivity (Shade & Handelsman, 2012; Busby *et al*., 2017; Toju *et al*., 2018). Despite being derived from a single soil source and wheat cultivar, TriMic captured 72% of genera consistently found in 50% of studies reviewed by Kavamura *et al*. (2021), including taxa previously identified as hub or keystone members of the wheat microbiome, such as *Sphingomonas, Telluria (Massilia)*, and *Luteibacter* (Araujo *et al*., 2019; Schlatter *et al*., 2020; Simonin *et al*., 2020). Hub taxa are thought to play important roles in community stability and function (Toju *et al*., 2018). The absence of the remaining core genera (*Bradyrhizobium, Chitinophaga*, and *Pedobacter*) and other wheat-associated taxa, such as

*Streptomyces,Nitrospira, Nitrosovibrio*, and *Nitrosospira*, may reflect cultivation biases, particularly for low-abundance or slow growing organisms that require specialized isolation strategies (Amann *et al*., 1995; Vartoukian *et al*., 2010; Pham & Kim, 2012). Additionally, the collection includes members of the genus *Kosakonia*, which to our knowledge, has not been previously reported as part of the wheat microbiome, but has been isolated from fava bean and shown to promote growth of wheat under high salinity (Ali & El-Kholy, 2022). Therefore, its inclusion expands the collection’s functional relevance beyond the core microbiota.

A limitation of focusing on highly prevalent and core taxa is that it may bias towards generalist organisms due to their metabolic plasticity and ability to occupy a wide range of niches in the plant environment rather than reflecting close associations with host function (Neu *et al*., 2021). This could result in overlooking rare or highly specialized microbes that contribute to host function under certain conditions. Even though isolation coverage in other plant associated collections has been defined based on full microbiome coverage or occupancy-abundance curves from culture independent studies (Zhalnina *et al*., 2018; Dai *et al*., 2025; Garrell *et al*., 2026b), these methods suffer from the same limitation as ecological function cannot be inferred from amplicon or even full-genome-based analyses. Regardless of coverage assessment method, the major limitation remains cultivation biases which limit the recovery of all taxa. Future efforts should therefore prioritize targeted recovery of functionally important groups missing from this collection and expand beyond the bacterial component of the microbiome to include fungi, archaea, and protists, which also play key roles in community assembly and plant performance (Gao *et al*., 2019; Jung *et al*., 2020; Murali *et al*., 2021); an undertaking to be facilitated by the DSMZ microbiome collection concept which allows for future expansion of TriMic while maintaining public accessibility.

Although the functional core microbiome has been proposed as a more comprehensive approach to determine ecological relevance of bacterial collections, its definition remains unresolved (Lemanceau *et al*., 2017). Therefore, the functional relevance of plant microbial collections has been assessed by determining the presence of target genes in similar approaches to the one describe here (Dai *et al*., 2025; Garrell *et al*., 2026b). Despite differences in isolation strategies, the plant culture collections produced by Dai *et al*. (2025) and Garrell *et al*. (2026b) also presented broad functional capabilities across taxonomic groups with functional profiles ranging from generalists to specialists. While none of the isolates encoded all target functions, this does not preclude their relevance, as plant colonization and host-associated processes are often supported by complementary traits distributed across multiple taxa rather than a single organism (do Amaral *et al*., 2020). Moreover, the prevalence of biosynthetic gene clusters associated with non-ribosomal peptides, terpenes, aryl polyenes, and homoserine lactones is consistent with patterns observed in other plant-associated genomes and culture collections (Mukherjee *et al*., 2023; Dai *et al*., 2025). Secondary metabolites support diverse roles in plant-microbe and microbe-microbe interactions (Pang *et al*., 2021; Ravelo-Ortega *et al*., 2023) including pathogen suppression and modulate microbe-microbe and plant-microbe interactions (Ullah *et al*., 2020; Sharifi *et al*., 2010; Zarkani *et al*., 2013; Hartmann *et al*., 2021).Overall, these findings demonstrates that a host-mediated approach can be effectively applied for microbial isolation, resulting in the recovery of major plant-associated taxa and functional traits relevant in plant-microbiota interactions. Although, genome-based screening provides valuable insights into functional potential, experimental validation is necessary to determine the conditions under which these traits are expressed *in vivo*. Importantly, the functional overview provided here extends the utility of the collection by guiding strain selection and enabling the identification of isolates with traits of interest for hypothesis-driven experimental design. Therefore, highlighting the utility of this resource to interrogate different aspects of plant-microbe and microbe-microbe interactions under different contexts. In this context, the combination of functional diversity and redundancy observed in this collection may enhance the resilience of SynComs derived from it under variable conditions.

Different strategies can be used to assemble SynComs (Xu *et al*., 2025), each tailored to target different aspects of the microbiome. We used a host-mediated approach to select members with taxonomic and functional traits relevant to mechanistic studies in plant-microbe interactions. The proposed 15-member SynCom incorporates core and putative keystone taxa associated with the wheat microbiome. Moreover, its predicted functional composition reflects that of the broader collection, maintaining functional redundancy and encompassing both generalist and specialist. Host-mediated selection also led to the development of a reduced maize SynCom that has allowed for the exploration of the ecological relevance of keystone taxa (Niu *et al*., 2017), heterosis effect (Wagner *et al*., 2021), and the identification of colonization functions expressed *in planta* through metaproteomics (Garrell *et al*., 2026a). These studies validate host-mediated approaches to generate tractable communities that retain ecologically relevant features of the microbiome. Nevertheless, SynComs can also be developed based on microbial functional traits known to benefit the host. This strategy has resulted in SynComs in wheat with fungal biocontrol function (Liu *et al*., 2022), and salt and drought stress tolerance (Khan *et al*., 2022; Rostamian *et al*., 2023). Another common SynCom building strategy is to select taxa that are highly abundant in the host microbiome. Using this approach a SynCom capable of the recapitulating the colonization patterns of a natural community was developed in wheat (Bak *et al*., 2026). Notably, all SynCom building approaches successfully achieved their intended objectives, suggesting that SynCom design is not defined by an optimal strategy, but rather its suitability for addressing a given research question by capturing the necessary taxonomic and functional traits (Northen *et al*., 2024).

At the same time, each assembly method carries inherent limitations. For example, host-mediated and functionally based selection may bias toward the environmental conditions under which selection occurs, potentially limiting performance under alternative contexts. More broadly, SynCom performance remains highly context-dependent, as community behavior may shift across environmental conditions. For this reason, the stability and reproducibility of the proposed wheat SynCom should be experimentally validated to evaluate its community dynamics across environmental conditions. Additionally, small SynComs such as the one presented here are not appropriate for analyses that rely on higher community complexity, such as co-occurrence network inference (Vorholt *et al*., 2017; Liu *et al*., 2019; Marín *et al*., 2021; Xu *et al*., 2025). Despite these limitations, the proposed SynCom was developed through an ecologically informed selection process based on observed community dynamics and provides a defined starting point for developing a tractable platform with a broad range of functional capabilities to interrogate microbial contributions to plant responses and community interactions.

Overall, TriMic expands the toolkit available for research on wheat-microbe interactions by providing a publicly accessible and versatile resource for hypothesis-driven research. By integrating insights from the literature with the functional overview of the collection, targeted hypotheses can be tested by selectively removing, replacing, or adding microbes or genes to the proposed SynCom. For example, removing putative keystone species could reveal their roles in community structure and function, while exchanging strains with contrasting functional traits can disentangle their contributions to community assembly and plant performance. Additionally, other isolates within the collection can be used to produce higher-complexity SynComs if needed. Moreover, the introduction of additional bacterial or fungal members could enable the study of invasion dynamics and their effects on plant health and community structure. Together, TriMic and the proposed SynCom provide a foundation for mechanistic studies of wheat-microbe interactions, with the potential advancement of our understanding of plant-microbe interactions and support the development of novel bioinoculants to enhance agricultural productivity.

## Acknowledgements

We thank all the curators and technicians of the German Collection of Microorganisms and Cell Cultures (DSMZ) for their contributions during preservation and authentication of the collection, and Ruthie Stokes for their assistance with experimental set up.

## Author contributions

AG, CSJ, MK, and PS acquired funding. CT, AG, and MK contributed to conceptualization and experimental design. CT led the study, performed experiments, bioinformatics, and statistical analysis, and wrote the original manuscript draft. AZ and CSJ provided supervision and resources for whole genome sequencing and bioinformatic analysis. AW assisted with experimental work. FN contributed by running AntiSMASH analysis and providing summaries under the supervision of PS. LEJ carried out amplicon sequencing. RH led and coordinated microbial strain curation and generation of microbiome collection at the DSMZ. All authors reviewed and edited the manuscript.

## Data availability

Sequencing data generated in this study are publicly available in the NCBI database under BioProject accession numbers PRJNA1147241 and PRJNA1415243.

## Supporting Information

Supplementary text. Expanded results and discussion of genome functional analysis and biosynthetic gene clusters.

Fig S1. Phylogenetic tree of 99 sequenced isolates based on 16S rRNA gene sequences.

Fig S2. Distribution of the top ten biosynthetic gene clusters (BGCs) identified across representative isolates (n=69) from each clade in the TriMic collection available at the DSMZ.

Fig S3. Mean wheat responses to consortium inoculation at 15 days post emergence (Mean ± standard error).

Fig S4. Relative read abundance (mean ±SE) of individual consortium strains in the wheat cultivar Catawba at 5, 10, and 15 days post-emergence (DPE).

Fig S5. Relative read abundance (mean ±SE) of individual consortium strains in across the four wheat genotypes at 15 days post-emergence (DPE).

Fig S6. Wheat root synthetic community (SynCom) functional traits.

## Supporting Data

Table S1. List of KEGG ortholog (KO) numbers of target genomic features associated with bacterial adaptations to the plant environment, nitrogen and phosphorus cycling, and plant growth promotion.

Table S2. Bacterial isolates metadata collected during isolation. Colony morphology descriptions are based on observations made on isolation media.

Table S3. Assigned genera identified in the original root slurry sample with percent relative read abundances.

Table S4. List of taxa associated with wheat bacterial microbiota as compiled by Kavamura et al. (2021) in Figure 2. Genera in bold are present in TriMic.

Supplementary Table 5. Quality control metrics of final genome assemblies. Data shown for both short read and hybrid assemblies.

Table S6. Isolate classification based on whole genome sequences assigned using the genome taxonomy database toolkit (GTDB-tk) and the type strain genome server (TYGS).

Table S7. Results of gene screen on DSMZ collection genomes (n=88). Gene present (0) Gene absent (1).

Table S8. Number of biosynthetic gene clusters identified in AntiSMASH by class found in representative strains of the DSMZ bacterial collection (n=69).

Table S9. List of identified metabolites potentially produced by representative strains of the DSMZ bacterial collection (n=69) based on AntiSMASH analysis.

Table S10. Two-way analysis of variance of wheat plant performance measurements at 15 days post-emergence. Significant effects are indicated in bold (P < 0.05)

Table S11. Comparison of consortium strain relative abundance in wheat roots at 15 days post emergence across all genotypes with the initial consortium inoculum using a one sample t-tests.

Table S12. DESeq2 pairwise contrasts of consortium strains abundance among 5, 10, and 15 days post-emergence (DPE).

Table S13. DESeq2 pairwise contrasts of consortium strains abundance among wheat genotypes.

## References

Ali Amal A, El-Kholy Aziza S. 2022. Isolation and Characterization of Endophytic Kosakonia radicincitans to Stimulate Wheat Growth in Saline Soil. Journal of Advances in Microbiology: 115–126.

Amann RI, Ludwig W, Schleifer KH. 1995. Phylogenetic identification and in situ detection of individual microbial cells without cultivation. Microbiological Reviews 59: 143–169.

do Amaral Fernanda Plucani, Tuleski Thalita Regina, Pankievicz Vania Carla Silva, Melnyk Ryan A, Arkin Adam P, Griffitts J, Tadra-Sfeir MZ, Maltempi de Souza E, Deutschbauer A, Monteiro RA, et al. 2020. Diverse Bacterial Genes Modulate Plant Root Association by Beneficial Bacteria. mBio 11: 10.1128/mbio.03078-20.

Araujo R, Dunlap C, Barnett S, Franco CMM. 2019. Decoding Wheat Endosphere–Rhizosphere Microbiomes in Rhizoctonia solani–Infested Soils Challenged by Streptomyces Biocontrol Agents. Frontiers in Plant Science 10: 1038.

Bai Y, Müller DB, Srinivas G, Garrido-Oter R, Potthoff E, Rott M, Dombrowski N, Münch PC, Spaepen S, Remus-Emsermann M, et al. 2015. Functional overlap of the Arabidopsis leaf and root microbiota. Nature 528: 364–369.

Bak F, Meier JK, Markussen B, Molina-Zamudio KY, Tang C, Nicolaisen MH. 2026. Synthetic bacterial community colonizes wheat roots grown in soil and mimics the assembly pattern of a field community in a cultivar dependent manner. ISME Communications 6: ycag028.

Bakker MG, McCormick SP. 2019. Microbial Correlates of Fusarium Load and Deoxynivalenol Content in Individual Wheat Kernels. Phytopathology® 109: 993–1002.

Bankevich A, Nurk S, Antipov D, Gurevich AA, Dvorkin M, Kulikov AS, Lesin VM, Nikolenko SI, Pham S, Prjibelski AD, et al. 2012. SPAdes: A New Genome Assembly Algorithm and Its Applications to Single-Cell Sequencing. Journal of Computational Biology 19: 455–477.

Batista BD, Singh BK. 2021. Realities and hopes in the application of microbial tools in agriculture. Microbial Biotechnology 14: 1258–1268.

Becker LE, Marshall D, Cubeta MA. 2024. A synergistic culture dependent and independent approach reveals a conserved wheat seed mycobiome.: 2024.02.22.581674.

Berendsen RL, Vismans G, Yu K, Song Y, de Jonge R, Burgman WP, Burmølle M, Herschend J, Bakker PAHM, Pieterse CMJ. 2018. Disease-induced assemblage of a plant-beneficial bacterial consortium. The ISME Journal 12: 1496–1507.

Bolyen E, Rideout JR, Dillon MR, Bokulich NA, Abnet CC, Al-Ghalith GA, Alexander H, Alm EJ, Arumugam M, Asnicar F, et al. 2019. Reproducible, interactive, scalable and extensible microbiome data science using QIIME 2. Nature Biotechnology 37: 852–857.

van den Boogaart KG, Tolosana-Delgado R, Bren M. 2025. compositions: Compositional Data Analysis.

Bruto M, Prigent-Combaret C, Muller D, Moënne-Loccoz Y. 2014. Analysis of genes contributing to plant-beneficial functions in plant growth-promoting rhizobacteria and related Proteobacteria. Scientific Reports 4: 6261.

Buchfink B, Reuter K, Drost H-G. 2021. Sensitive protein alignments at tree-of-life scale using DIAMOND. Nature Methods 18: 366–368.

Busby PE, Soman C, Wagner MR, Friesen ML, Kremer J, Bennett A, Morsy M, Eisen JA, Leach JE, Dangl JL. 2017. Research priorities for harnessing plant microbiomes in sustainable agriculture. PLOS Biology 15: e2001793.

Bushnell B. 2014. BBtools.

Callahan BJ, McMurdie PJ, Rosen MJ, Han AW, Johnson AJA, Holmes SP. 2016. DADA2: High-resolution sample inference from Illumina amplicon data. Nature Methods 13: 581–583.

Cantalapiedra CP, Hernández-Plaza A, Letunic I, Bork P, Huerta-Cepas J. 2021. eggNOG-mapper v2: Functional Annotation, Orthology Assignments, and Domain Prediction at the Metagenomic Scale. Molecular Biology and Evolution 38: 5825–5829.

Carlström CI, Field CM, Bortfeld-Miller M, Müller B, Sunagawa S, Vorholt JA. 2019. Synthetic microbiota reveal priority effects and keystone strains in the Arabidopsis phyllosphere. Nature Ecology & Evolution 3: 1445–1454.

Castrillo G, Teixeira PJPL, Paredes SH, Law TF, de Lorenzo L, Feltcher ME, Finkel OM, Breakfield NW, Mieczkowski P, Jones CD, et al. 2017. Root microbiota drive direct integration of phosphate stress and immunity. Nature 543: 513–518.

Chaumeil P-A, Mussig AJ, Hugenholtz P, Parks DH. 2020. GTDB-Tk: a toolkit to classify genomes with the Genome Taxonomy Database. Bioinformatics 36: 1925–1927.

Chen S, Waghmode TR, Sun R, Kuramae EE, Hu C, Liu B. 2019. Root-associated microbiomes of wheat under the combined effect of plant development and nitrogen fertilization. Microbiome 7: 136.

Chin-A-Woeng TFC, Bloemberg GV, Mulders IHM, Dekkers LC, Lugtenberg BJJ. 2000. Root Colonization by Phenazine-1-Carboxamide-Producing Bacterium Pseudomonas chlororaphis PCL1391 Is Essential for Biocontrol of Tomato Foot and Root Rot. Molecular Plant-Microbe Interactions® 13: 1340–1345.

Cole BJ, Feltcher ME, Waters RJ, Wetmore KM, Mucyn TS, Ryan EM, Wang G, Ul-Hasan S, McDonald M, Yoshikuni Y, et al. 2017. Genome-wide identification of bacterial plant colonization genes. PLOS Biology 15: e2002860.

Collyer ML, Adams DC. 2018. RRPP: An r package for fitting linear models to high-dimensional data using residual randomization. Methods in Ecology and Evolution 9: 1772–1779.

Compant S, Clément C, Sessitsch A. 2010. Plant growth-promoting bacteria in the rhizo- and endosphere of plants: Their role, colonization, mechanisms involved and prospects for utilization. Soil Biology and Biochemistry 42: 669–678.

Dai R, Zhang J, Liu F, Xu H, Qian J-M, Cheskis S, Liu W, Wang B, Zhu H, Pronk LJU, et al. 2025. Crop root bacterial and viral genomes reveal unexplored species and microbiome patterns. Cell 188: 2521–2539.e22.

Dastogeer KMG, Tumpa FH, Sultana A, Akter MA, Chakraborty A. 2020. Plant microbiome–an account of the factors that shape community composition and diversity. Current Plant Biology 23: 100161.

Durán P, Thiergart T, Garrido-Oter R, Agler M, Kemen E, Schulze-Lefert P, Hacquard S. 2018. Microbial Interkingdom Interactions in Roots Promote Arabidopsis Survival. Cell 175: 973–983.e14.

Erenstein O, Jaleta M, Mottaleb KA, Sonder K, Donovan J, Braun H-J. 2022. Global Trends in Wheat Production, Consumption and Trade. In: Reynolds MP, Braun H-J, eds. Wheat Improvement: Food Security in a Changing Climate. Cham: Springer International Publishing, 47–66.

Forry SP, Servetas SL, Kralj JG, Soh K, Hadjithomas M, Cano R, Carlin M, de Amorim MG, Auch B, Bakker MG, et al. 2024. Variability and bias in microbiome metagenomic sequencing: an interlaboratory study comparing experimental protocols. Scientific Reports 14: 9785.

Frøslev TG, Kjøller R, Bruun HH, Ejrnæs R, Brunbjerg AK, Pietroni C, Hansen AJ. 2017. Algorithm for post-clustering curation of DNA amplicon data yields reliable biodiversity estimates. Nature Communications 8: 1188.

Fu L, Niu B, Zhu Z, Wu S, Li W. 2012. CD-HIT: accelerated for clustering the next-generation sequencing data. Bioinformatics 28: 3150–3152.

Gao Z, Karlsson I, Geisen S, Kowalchuk G, Jousset A. 2019. Protists: Puppet Masters of the Rhizosphere Microbiome. Trends in Plant Science 24: 165–176.

Garrell A-K, Cheadle J, Crook N, Pal G, Septer AN, Wagner MR, Beck AE, Kleiner M. 2026a. Differential proteomics of bacteria grown in vitro and in planta reveals functions used during growth on maize roots. mSystems 11: e0037126.

Garrell A-K, Ginnan N, Swift JF, Pal G, Zervas A, Pestalozzi C, Tang C, Tso F, Ford NE, Niu B, et al. 2026b. ZeaMiC: a Publicly Available Culture Collection of Maize Root-Associated Bacteria.: 2026.03.23.713778.

Garrido-Sanz D, Keel C. 2025. Seed-borne bacteria drive wheat rhizosphere microbiome assembly via niche partitioning and facilitation. Nature Microbiology 10: 1130–1144.

Gdanetz K, Trail F. 2017. The Wheat Microbiome Under Four Management Strategies, and Potential for Endophytes in Disease Protection. Phytobiomes Journal 1: 158–168.

Hansen CHF, Krych L, Nielsen DS, Vogensen FK, Hansen LH, Sørensen SJ, Buschard K, Hansen AK. 2012. Early life treatment with vancomycin propagates Akkermansia muciniphila and reduces diabetes incidence in the NOD mouse. Diabetologia 55: 2285–2294.

Hartmann A, Klink S, Rothballer M. 2021. Importance of N-Acyl-Homoserine Lactone-Based Quorum Sensing and Quorum Quenching in Pathogen Control and Plant Growth Promotion. Pathogens 10: 1561.

Hu J, Fan J, Sun Z, Liu S. 2020. NextPolish: a fast and efficient genome polishing tool for long-read assembly. Bioinformatics 36: 2253–2255.

Johnston-Monje D, Gutiérrez JP, Lopez-Lavalle LAB. 2021. Seed-Transmitted Bacteria and Fungi Dominate Juvenile Plant Microbiomes. Frontiers in Microbiology 12.

Jung J, Kim J-S, Taffner J, Berg G, Ryu C-M. 2020. Archaea, tiny helpers of land plants. Computational and Structural Biotechnology Journal 18: 2494–2500.

Kanehisa M, Sato Y, Kawashima M, Furumichi M, Tanabe M. 2016. KEGG as a reference resource for gene and protein annotation. Nucleic Acids Research 44: D457–D462.

Kavamura VN, Mendes R, Bargaz A, Mauchline TH. 2021. Defining the wheat microbiome: Towards microbiome-facilitated crop production. Computational and Structural Biotechnology Journal 19: 1200–1213.

Kavamura VN, Robinson RJ, Hayat R, Clark IM, Hughes D, Rossmann M, Hirsch PR, Mendes R, Mauchline TH. 2019. Land Management and Microbial Seed Load Effect on Rhizosphere and Endosphere Bacterial Community Assembly in Wheat. Frontiers in Microbiology 10.

Khan MY, Nadeem SM, Sohaib M, Waqas MR, Alotaibi F, Ali L, Zahir ZA, Al-Barakah FNI. 2022. Potential of plant growth promoting bacterial consortium for improving the growth and yield of wheat under saline conditions. Frontiers in Microbiology 13: 958522.

Knights HE, Ramachandran VK, Jorrin B, Ledermann R, Parsons JD, Aroney STN, Poole PS. 2024. Rhizobium determinants of rhizosphere persistence and root colonization. The ISME Journal 18: wrae072.

Kolmogorov M, Yuan J, Lin Y, Pevzner PA. 2019. Assembly of long, error-prone reads using repeat graphs. Nature Biotechnology 37: 540–546.

Krueger F. 2021. Trim Galore!

Kwak M-J, Kong HG, Choi K, Kwon S-K, Song JY, Lee J, Lee PA, Choi SY, Seo M, Lee HJ, et al. 2018. Rhizosphere microbiome structure alters to enable wilt resistance in tomato. Nature Biotechnology 36: 1100–1109.

Lakshmanan V, Ray P, Craven KD. 2017. Rhizosphere Sampling Protocols for Microbiome (16S/18S/ITS rRNA) Library Preparation and Enrichment for the Isolation of Drought Tolerance-Promoting Microbes. In: Sunkar R, ed. Methods in Molecular Biology. Plant Stress Tolerance: Methods and Protocols. New York, NY: Springer, 349–362.

Lazcano C, Boyd E, Holmes G, Hewavitharana S, Pasulka A, Ivors K. 2021. The rhizosphere microbiome plays a role in the resistance to soil-borne pathogens and nutrient uptake of strawberry cultivars under field conditions. Scientific Reports 11: 3188.

Lemanceau P, Blouin M, Muller D, Moënne-Loccoz Y. 2017. Let the Core Microbiota Be Functional. Trends in Plant Science 22: 583–595.

Lenth RV, Piaskowski J. 2025. emmeans: Estimated Marginal Means, aka Least-Squares Means.

Letunic I, Bork P. 2024. Interactive Tree of Life (iTOL) v6: recent updates to the phylogenetic tree display and annotation tool. Nucleic Acids Research 52: W78–W82.

Levy A, Salas Gonzalez I, Mittelviefhaus M, Clingenpeel S, Herrera Paredes S, Miao J, Wang K, Devescovi G, Stillman K, Monteiro F, et al. 2018. Genomic features of bacterial adaptation toplants. Nature Genetics 50: 138–150.

Li J, Wang J, Liu H, Macdonald CA, Singh BK. 2022. Application of microbial inoculants significantly enhances crop productivity: A meta-analysis of studies from 2010 to 2020. Journal of Sustainable Agriculture and Environment 1: 216–225.

Liu Y-X, Qin Y, Bai Y. 2019. Reductionist synthetic community approaches in root microbiome research. Current Opinion in Microbiology 49: 97–102.

Liu H, Qiu Z, Ye J, Verma JP, Li J, Singh BK. 2022. Effective colonisation by a bacterial synthetic community promotes plant growth and alters soil microbial community. Journal of Sustainable Agriculture and Environment 1: 30–42.

Love MI, Huber W, Anders S. 2014. Moderated estimation of fold change and dispersion for RNA-seq data with DESeq2. Genome Biology 15: 550.

Marín O, González B, Poupin MJ. 2021. From Microbial Dynamics to Functionality in the Rhizosphere: A Systematic Review of the Opportunities With Synthetic Microbial Communities. Frontiers in Plant Science 12.

Mavrodi DV, Mavrodi OV, Elbourne LDH, Tetu S, Bonsall RF, Parejko J, Yang M, Paulsen IT, Weller DM, Thomashow LS. 2018. Long-Term Irrigation Affects the Dynamics and Activity of the Wheat Rhizosphere Microbiome. Frontiers in Plant Science 9.

McMurdie PJ, Holmes S. 2013. phyloseq: An R Package for Reproducible Interactive Analysis and Graphics of Microbiome Census Data. PLOS ONE 8: e61217.

Medema MH, Blin K, Cimermancic P, de Jager V, Zakrzewski P, Fischbach MA, Weber T, Takano E, Breitling R. 2011. antiSMASH: rapid identification, annotation and analysis of secondary metabolite biosynthesis gene clusters in bacterial and fungal genome sequences. Nucleic Acids Research 39: W339–W346.

Mehlferber EC, Arnault G, Joshi B, Partida-Martinez LP, Patras KA, Simonin M, Koskella B. 2024. A cross-systems primer for synthetic microbial communities. Nature Microbiology 9: 2765–2773.

Meier-Kolthoff JP, Göker M. 2019. TYGS is an automated high-throughput platform for state-of-the-art genome-based taxonomy. Nature Communications 10: 2182.

Mesny F, Hacquard S, Thomma BP. 2023. Co-evolution within the plant holobiont drives host performance. EMBO reports 24: e57455.

Mitter EK, Tosi M, Obregón D, Dunfield KE, Germida JJ. 2021. Rethinking Crop Nutrition in Times of Modern Microbiology: Innovative Biofertilizer Technologies. Frontiers in Sustainable Food Systems 5.

Mukherjee A, Tikariha H, Bandla A, Pavagadhi S, Swarup S. 2023. Global analyses of biosynthetic gene clusters in phytobiomes reveal strong phylogenetic conservation of terpenes and aryl polyenes. mSystems 8: e00387–23.

Murali M, Naziya B, Ansari MA, Alomary MN, AlYahya S, Almatroudi A, Thriveni MC, Gowtham HG, Singh SB, Aiyaz M, et al. 2021. Bioprospecting of Rhizosphere-Resident Fungi: Their Role and Importance in Sustainable Agriculture. Journal of Fungi 7: 314.

Neu AT, Allen EE, Roy K. 2021. Defining and quantifying the core microbiome: Challenges and prospects. Proceedings of the National Academy of Sciences 118: e2104429118.

Niu B, Paulson JN, Zheng X, Kolter R. 2017. Simplified and representative bacterial community of maize roots. Proceedings of the National Academy of Sciences 114: E2450–E2459.

Northen TR, Kleiner M, Torres M, Kovács ÁT, Nicolaisen MH, Krzyżanowska DM, Sharma S, Lund G, Jelsbak L, Baars O, et al. 2024. Community standards and future opportunities for synthetic communities in plant–microbiota research. Nature Microbiology 9: 2774–2784.

Ofek-Lalzar M, Sela N, Goldman-Voronov M, Green SJ, Hadar Y, Minz D. 2014. Niche and host-associated functional signatures of the root surface microbiome. Nature Communications 5: 4950.

Oksanen J, Simpson GL, Blanchet FG, Kindt R, Legendre P, Minchin PR, O’Hara RB, Solymos P, Stevens MHH, Szoecs E, et al. 2025. vegan: Community Ecology Package.

Olm MR, Brown CT, Brooks B, Banfield JF. 2017. dRep: a tool for fast and accurate genomic comparisons that enables improved genome recovery from metagenomes through de-replication. The ISME Journal 11: 2864–2868.

Pagé AP, Tremblay J, Masson L, Greer CW. 2019. Nitrogen- and phosphorus-starved Triticum aestivum show distinct belowground microbiome profiles (A Zhang, Ed.). PLOS ONE 14: e0210538.

Pang Z, Chen J, Wang T, Gao C, Li Z, Guo L, Xu J, Cheng Y. 2021. Linking Plant Secondary Metabolites and Plant Microbiomes: A Review. Frontiers in Plant Science 12.

Parks DH, Imelfort M, Skennerton CT, Hugenholtz P, Tyson GW. 2015. CheckM: assessing the quality of microbial genomes recovered from isolates, single cells, and metagenomes. Genome Research 25: 1043–1055.

Pham VHT, Kim J. 2012. Cultivation of unculturable soil bacteria. Trends in Biotechnology 30: 475–484.

Price MN, Dehal PS, Arkin AP. 2010. FastTree 2 – Approximately Maximum-Likelihood Trees for Large Alignments. PLOS ONE 5: e9490.

Pruesse E, Peplies J, Glöckner FO. 2012. SINA: Accurate high-throughput multiple sequence alignment of ribosomal RNA genes. Bioinformatics 28: 1823–1829.

Rascovan N, Carbonetto B, Perrig D, Díaz M, Canciani W, Abalo M, Alloati J, González-Anta G, Vazquez MP. 2016. Integrated analysis of root microbiomes of soybean and wheat from agricultural fields. Scientific Reports 6: 28084.

Ravelo-Ortega G, Raya-González J, López-Bucio J. 2023. Compounds from rhizosphere microbes that promote plant growth. Current Opinion in Plant Biology 73: 102336.

Rigerte L, Heintz-Buschart A, Reitz T, Tarkka MT. 2025. Assembly and application of a synthetic bacterial community for enhancing barley tolerance to drought. Frontiers in Bacteriology 4: 1572294.

Risely A. 2020. Applying the core microbiome to understand host–microbe systems. Journal of Animal Ecology 89: 1549–1558.

Robertson-Albertyn S, Abbott JC, Concas F, Brown LH, Orr JN, George TS, Bulgarelli D. 2022. Genome-Annotated Bacterial Collection of the Barley Rhizosphere Microbiota. Microbiology Resource Announcements 11: e01064–21.

Rossmann M, Pérez-Jaramillo JE, Kavamura VN, Chiaramonte JB, Dumack K, Fiore-Donno AM, Mendes LW, Ferreira MMC, Bonkowski M, Raaijmakers JM, et al. 2020. Multitrophic interactions in the rhizosphere microbiome of wheat: from bacteria and fungi to protists. FEMS Microbiology Ecology 96: fiaa032.

Rostamian A, Moaveni P, MehdiSadeghi-Shoae, Mozafari H, Rajabzadeh F. 2023. Effective drought mitigation by rhizobacteria consortium in wheat field trials. Rhizosphere 25: 100653.

Schlatter DC, Yin C, Hulbert S, Paulitz TC. 2020. Core Rhizosphere Microbiomes of Dryland Wheat Are Influenced by Location and Land Use History. Applied and Environmental Microbiology 86: e02135–19.

Seemann T. 2015. Snippy: fast bacterial variant calling from NGS reads.

Selten G, Lamouche F, Gómez-Repollés A, López JL, Zhang X, Smart C, Blahovska Z, Zarate Camargo G, Kelly S, de Jonge R, et al. 2026. Functional capacities drive recruitment of bacteria into plant root microbiota. Nature Microbiology 11: 2994–3009.

Seppey M, Manni M, Zdobnov EM. 2019. BUSCO: Assessing Genome Assembly and Annotation Completeness. In: Kollmar M, ed. Methods in Molecular Biology. Gene Prediction: Methods and Protocols. New York, NY: Springer, 227–245.

Shade A, Handelsman J. 2012. Beyond the Venn diagram: the hunt for a core microbiome. Environmental Microbiology 14: 4–12.

Sharifi R, Sharifi-Tehrani A, Talebi-Jahromi K, Ahmadzadeh M. 2010. Pyoverdine Production in Pseudomonas Fluorescens UTPF5 and its Association with Suppression of Common Bean Damping off Caused by Rhizoctonia Solani (Kühn). Journal of Plant Protection Research; 2010; *vol.* 50*; No* 1.

Shiferaw B, Smale M, Braun H-J, Duveiller E, Reynolds M, Muricho G. 2013. Crops that feed the world 10. Past successes and future challenges to the role played by wheat in global food security. Food Security 5: 291–317.

Simonin M, Dasilva C, Terzi V, Ngonkeu ELM, Diouf D, Kane A, Béna G, Moulin L. 2020. Influence of plant genotype and soil on the wheat rhizosphere microbiome: Evidences for a core microbiome across eight African and European soils. FEMS Microbiology Ecology 96.

Toju H, Peay KG, Yamamichi M, Narisawa K, Hiruma K, Naito K, Fukuda S, Ushio M, Nakaoka S, Onoda Y, et al. 2018. Core microbiomes for sustainable agroecosystems. Nature Plants 4: 247–257.

Trivedi P, Leach JE, Tringe SG, Sa T, Singh BK. 2020. Plant–microbiome interactions: from community assembly to plant health. Nature Reviews Microbiology 18: 607–621.

Tu Q, Lin L, Cheng L, Deng Y, He Z. 2019. NCycDB: a curated integrative database for fast and accurate metagenomic profiling of nitrogen cycling genes. Bioinformatics 35: 1040–1048.

Ullah A, Bano A, Janjua HT. 2020. Chapter 3 - Microbial Secondary Metabolites and Defense of Plant Stress. In: Singh JS, Vimal SR, eds. Microbial Services in Restoration Ecology. Elsevier, 37–46.

Vannier N, Mesny F, Getzke F, Chesneau G, Dethier L, Ordon J, Thiergart T, Hacquard S. 2023. Genome-resolved metatranscriptomics reveals conserved root colonization determinants in a synthetic microbiota. Nature Communications 14: 8274.

Vartoukian SR, Palmer RM, Wade WG. 2010. Strategies for culture of ‘unculturable’ bacteria. FEMS Microbiology Letters 309: 1–7.

Verma SK, Kharwar RN, Gond SK, Kingsley KL, White JF. 2019. Exploring Endophytic Communities of Plants: Methods for Assessing Diversity, Effects on Host Development and Potential Biotechnological Applications. In: Verma SK, White J James Francis, eds. Seed Endophytes: Biology and Biotechnology. Cham: Springer International Publishing, 55–82.

Vorholt JA, Vogel C, Carlström CI, Müller DB. 2017. Establishing Causality: Opportunities of Synthetic Communities for Plant Microbiome Research. Cell Host & Microbe 22: 142–155.

Wagner MR, Tang C, Salvato F, Clouse KM, Bartlett A, Vintila S, Phillips L, Sermons S, Hoffmann M, Balint-Kurti PJ, et al. 2021. Microbe-dependent heterosis in maize. Proceedings of the National Academy of Sciences 118: e2021965118.

Wick RR, Judd LM, Gorrie CL, Holt KE. 2017. Unicycler: Resolving bacterial genome assemblies from short and long sequencing reads. PLOS Computational Biology 13: e1005595.

Wick RR, Schultz MB, Zobel J, Holt KE. 2015. Bandage: interactive visualization of de novo genome assemblies. Bioinformatics 31: 3350–3352.

Wickham H. 2016. ggplot2: Elegant Graphics for Data Analysis. Springer-Verlag New York.

Wippel K, Tao K, Niu Y, Zgadzaj R, Guan R, Dahms E, Zhang P, Jensen DB, Logemann E, Radutoiu S, et al. 2021. Host preference and invasiveness of commensals in the Lotus and Arabidopsis root microbiota. bioRxiv: 2021.01.12.426357.

Xu X, Dinesen C, Pioppi A, Kovács ÁT, Lozano-Andrade CN. 2025. Composing a microbial symphony: synthetic communities for promoting plant growth. Trends in Microbiology 0.

Yin C, Hulbert SH, Schroeder KL, Mavrodi O, Mavrodi D, Dhingra A, Schillinger WF, Paulitz TC. 2013. Role of Bacterial Communities in the Natural Suppression of Rhizoctonia solani Bare Patch Disease of Wheat (Triticum aestivum L.). Applied and Environmental Microbiology 79: 7428–7438.

Zarkani AA, Stein E, Röhrich CR, Schikora M, Evguenieva-Hackenberg E, Degenkolb T, Vilcinskas A, Klug G, Kogel K-H, Schikora A. 2013. Homoserine Lactones Influence the Reaction of Plants to Rhizobia. International Journal of Molecular Sciences 14: 17122–17146.

Zeng J, Tu Q, Yu X, Qian L, Wang C, Shu L, Liu F, Liu S, Huang Z, He J, et al. 2022. PCycDB: a comprehensive and accurate database for fast analysis of phosphorus cycling genes. Microbiome 10: 101.

Zhalnina K, Louie KB, Hao Z, Mansoori N, da Rocha UN, Shi S, Cho H, Karaoz U, Loqué D, Bowen BP, et al. 2018. Dynamic root exudate chemistry and microbial substrate preferences drive patterns in rhizosphere microbial community assembly. Nature Microbiology 3: 470–480.

Zhao C, Liu B, Piao S, Wang X, Lobell DB, Huang Y, Huang M, Yao Y, Bassu S, Ciais P, et al. 2017. Temperature increase reduces global yields of major crops in four independent estimates. Proceedings of the National Academy of Sciences 114: 9326–9331.

