## Supplementary information for "TriMic: a *Triticum aestivum* microbial culture collection and synthetic community for dissecting wheat-microbe interactions"

Supporint Information for TriMic: a ***Tri****ticum aestivum* **mic**robial culture collection and synthetic community for dissecting wheat-microbe interactions

Clara Tang, Athanasios Zervas, Andrea Ward, Fani Ntana, Richard L. Hahnke, Lea Ellegaard-Jensen, Peter Stougaard, Carsten Suhr Jacobsen, Amy Grunden, Manuel Kleiner

Corresponding authors: Manuel Kleiner and Clara Tang

**File contains:**

Supporting Text

Figures S1-S6

### Supporting Text

*Distribution of functional traits indicates both generalist and specialist are present in the collection*

Genome screened revealed broad functional diversity across isolates, indicating a continuum of functional profiles within the collection. At the same time, substantial functional redundancy and complementarity were observed across phylogenetic groups, highlighting the collection a robust resource for the plant-microbiome studies (Northen *et al.*, 2024). Genes associated with plant growth promotion were found in 72 % of isolates in the collection, although their distribution varied markedly across taxa. The plant hormone regulator amino-cyclopropane carboxylate deaminase (ACC-deaminase) was restricted to *Pseudomonas* strains and members of the class Bacteroidia. In contrast, genes involved in auxin (indole-3- acetic acid, IAA) production were primarily identified in isolates within the *Enterobacteriaceae* family, with a few additional occurrences in strains Paensp111, Golu5, and Perisp51. Similarly, hydrogen cyanide (HCN) production genes were largely confined to members of the *Burkholderiaceae* family and *Pseudomonas* strains. In comparison, 2,3-butanediol synthesis was detected in only 23 strains in the collection, predominantly within the Actinomycetales order and the Firmicutes phylum. Strains carrying these functions could serve as bioinoculants because strains carrying such functions have been shown to enhance wheat growth under optimal and stress growth conditions (Chandra *et al.*, 2018; Emami *et al.*, 2019; Farahat *et al.*, 2020). However, the actual ability of these strains to perform these beneficial functions needs to be tested *in planta* and under different conditions for a better understanding of their potential implementation as bioinoculants.

Moreover, the collection also has potential for use as biofertilizers as nutrient cycling functions are also widespread. We observed that genes for the metabolism and transport of nitrogen (N) and phosphorus (P) were also highly prevalent in the collection, although variation emerged at the level of more specific functions. Transport of nitrate ($NO_3^-$) and nitrite ($NO_2^-$) was present in 89 % of the genomes, whereas urea transport was less common (69 %). Members of the *Curtobacterium* genus and strain Lutesp34 lack these N compound transport genes. Among N transformation pathways, nitrate reduction and denitrification genes were the most widespread, while nitrogen-fixing (*nif*HDK) genes were restricted to only three strains, Azosp132, Paensp38, and Pseusp138. Nitrogen-fixing bacteria within these genera have been shown to increase wheat yield under reduced N fertilization regimes in field conditions (Li *et al.*, 2019; Din *et al.*, 2021; Ebrahimi *et al.*, 2023). Ammonia oxidation genes were absent from all genome, revealing a functional gap in the collection and highlighting the need for targeted isolations efforts.

Phosphorus mobilization functions showed a similar pattern. While phosphate ($PO_4^{-2}$) transport genes were present in all genomes, transport systems for organic P (orgP) compounds were less prevalent. Glycerol phosphate and phosphonate transport genes were detected in approximately 75 % of the genomes; a higher proportion of these genes was found in the Rhizobiales order but were completely absent in members of the *Sphingomonas* genus and Bacteroidia class. Inositol transport genes were present in 24 % of genomes, primarily within the Rhizobiales order, some *Pseudomonas* strains, and isolates Padi120 and Burksp114. Genes involved in inorganic phosphorus (iP) mobilization were also widely distributed; genera with the highest number of genes in this category included *Massilia, Telluria, Rugamonas, Burkholderia, Paraburkholderia, Ralstonia*, and *Pseudomonas*, as well as strains Varisp41 and Rhizsp130, whereas these genes were absent in Golu5 and Perisp51. Similarly, orgP mobilization genes were present across the collection, with the exception of *Curtobacterium* strains, but generally at lower proportions. The greatest number of orgP mobilization genes was found in the Rhizobiales order and strain Perisp51. Similarly, wheat inoculated with P mobilizing bacteria improved growth and yield of wheat even in the absence of P fertilizers (Li *et al.*, 2020; Aziz *et al.*, 2020; Khourchi *et al.*, 2022; Wang *et al.*, 2022). Beyond their potential use as biofertilizers, the contrasting patterns of N and P transport and mobilization provide an opportunity to investigate the physiological basis of microbe-microbe interactions that maintain beneficial functions within synthetic communities (SynCom).

Genes associated with adaptations to plants and plant colonization were also widely distributed in the collection indicating ability to colonize the plant environment (Selten *et al.*, 2026). Genes involved in the metabolism and transport of amino acids and nucleotides, biofilm biosynthesis, quorum sensing, and metabolism of lipids, cofactors and vitamins were present in all genomes. Exopolysaccharide biosynthesis genes were found in 93 % of the collection, missing from strain Golu5, members of *Sphingomonas* genus, and the Xanthomonadales order. Sulfur (S) compound transport and relay system genes were present in approximately 85 % of genomes and in high proportion, although they were absent from most members of the Bacteroidia class and Xanthomonadales and Actinomycetales orders. Assimilatory sulfate reduction genes were present in 97 % of genomes, often with more than 50 % of genes in this category present; however, these genes were absent in *Curtobacterium* strains. In contrast, dissimilatory sulfate reduction was restricted to Dugasp25 and members of the *Massilia, Telluria, Burkholderia, Paraburkholderia,* and *Ralstonia* genera. Similarly, sulfate oxidation genes were limited to strains Golu5, Chor24, and Azosp132, as well as the members of *Duganella, Rugamonas, Burkholderia, Paraburkholderia, Ralstonia, Polaromonas, Mucilaginibacter*, and *Paenibacillus* genera. Genes associated with cell attachment and motility were also common. Pilus formation genes were present in 90 % of genomes, whereas fimbriae genes were found only in 58 %, mainly within members of the γ-Proteobacteria. Most strains encoded some form of motility, with chemotaxis being the most prevalent; only members of the *Micrococcaceae* family lacked motility-related genes. Finally, type I and VI bacterial secretion system genes were found in an average of 74 % of genomes, whereas Type III and IV were present in only 39 %. Most of the secretion systems were found within members of the Proteobacteria phylum, although some type I and VI secretion systems were also identified in members of the Bacteroidia class. Additionally, strains Perisp51 and Prme7 also contained type IV secretion system genes.  Several of these functions have been shown to increase in expression in the presence of plants (Garrell *et al.*, 2026), while their contrasting distribution across taxa suggests diverse strategies for colonizing different niches within the plant environment (do Amaral *et al.*, 2020). These constrasting functional profiles could also be leveraged in SynCom studies investigate potential cooperative interactions in root colonization (Selten *et al.*, 2026).

*Diverse biosynthetic gene clusters are distributed throughout the collection and show phylogenetic patterns*

In addition to screening for bacterial genes associated with plant interactions, we assessed the secondary metabolite production potential of the collection isolates by identifying biosynthetic gene clusters (BGCs). A total of 568 BGCs were identified in the collection, which were classified into 60 cluster classes (Table S8). The fewest BGCs were identified in *Arthrobacter* (Pseusp20), which only had three BGCs. In contrast, strains Paensp38 (*Paenibacillus)* and Azosp132 (*Azospirillum*) carried the highest number of BGCs, with 19 and 15 clusters identified, respectively. Based on the total number of clusters per class identified in the collection, the top ten classes identified were terpene production, unspecified ribosomally synthesized and post-translationally modified peptide products (RiPP-like), aryl polyenes, non-ribosomal peptide synthases (NRPS), redox cofactors, siderophores, β-lactones, type III polyketide synthase (T3PKS), and homoserine lactones (Fig S2). Secondary metabolites mediate plant-microbe and microbe-microbe interactions that can determine plant performance under certain conditions (Pang *et al.*, 2021; Ravelo-Ortega *et al.*, 2023).

The BGC patterns of different cluster classes differed based on taxonomy with a few noteworthy examples associated with metabolites known to be important for plant-microbe interactions. Gene clusters for the synthesis of siderophores such as bacillibactin, pyoverdin, crochelin A, graminibactin, malleobactin, xanthoferrin, ornibactin, and staphilloferrin were identified within the collection (Table  S9). However, these gene clusters were missing from strains in the *Xanthomonadales* order, *Enterobacteriaceae* family, and in genera *Variovorax, Paenibacillus, Mucilaginibacter, Azospirillum, Pseudarthrobacter*, and *Sphingomonas*. Siderophores are lowmolecular-weight compounds that bind insoluble iron in the environment and can suppress pathogen populations by limiting iron availability. Purified pyoverdine, desferal, and bacillibactin produced by bacteria were found to inhibit fungal pathogens (Sharifi *et al.*, 2010; Kumar *et al.*, 2024) and *Pseudomonas syringae* pv. Tomato (Dimopoulou *et al.*, 2021). Furthermore, many antimicrobial and cytotoxic products have been classified as NRPs, polyketides, terpenes, and ribosomal peptides (Ullah *et al.*, 2020). Antimicrobial compounds such as bacillomycin (Gu *et al.*, 2017), bottromycin(Franz *et al.*, 2021), fengycin (Ding *et al.*, 2025), formicamycins (Qin *et al.*, 2017), and polymyxin (Niu *et al.*, 2013) were identified within our isolates, thus demonstrating the great biocontrol potential of the collection.

Notably, the *Rhizobiaceae* family had the most homoserine lactone (AHL) BGCs identified. Interestingly, AHL lactone production was not prevalent in the collection, with only 14% of genomes containing BGCs for this type of molecule used in bacterial quorum sensing. Moreover, these molecules can also be recognized by plants and evoke specific outcomes. For example, *Arabidopsis thaliana* only showed increased resistance to pathogenic bacteria when inoculated with a long-chain AHL producing bacterial strain and not when inoculated a short-chain AHL producing strain (Zarkani *et al.*, 2013). Moreover, some AHL can regulate pathogen populations by quorum quenching and quorum sensing inhibition (Hartmann *et al.*, 2021). In addition, genes encoding specific linear and cyclic lipopeptides, pigments, and other antimicrobial and bioactive compounds were identified within throughout the collection, further highlighting its potential as a resource for studying pathogen suppression and plant-microbe interactions.

### References


do Amaral FP, Tuleski TR, Pankievicz VCS, Melnyk RA, Arkin AP, Griffitts J, Tadra-Sfeir MZ, Maltempi de Souza E, Deutschbauer A, Monteiro RA, *et al.* 2020. Diverse Bacterial Genes Modulate Plant Root Association by Beneficial Bacteria. *mBio* 11: 10.1128/mbio.03078-20.

Aziz MZ, Yaseen M, Naveed M, Wang X, Fatima K, Saeed Q, Mustafa A. 2020. Polymer-Paraburkholderia phytofirmans PsJN Coated Diammonium Phosphate Enhanced Microbial Survival, Phosphorous Use Efficiency, and Production of Wheat. *Agronomy* 10: 1344.

Chandra D, Srivastava R, Sharma AK. 2018. Influence of IAA and ACC Deaminase Producing Fluorescent Pseudomonads in Alleviating Drought Stress in Wheat (Triticum aestivum). *Agricultural Research* 7: 290–299.

Dimopoulou A, Theologidis I, Benaki D, Koukounia M, Zervakou A, Tzima A, Diallinas G, Hatzinikolaou DG, Skandalis N. 2021. Direct Antibiotic Activity of Bacillibactin Broadens the Biocontrol Range of Bacillus amyloliquefaciens MBI600. *mSphere* 6: 10.1128/msphere.00376-21.

Din I, Khan H, Ahmad Khan N, Khil A. 2021. Inoculation of nitrogen fixing bacteria in conjugation with integrated nitrogen sources induced changes in phenology, growth, nitrogen assimilation and productivity of wheat crop. *Journal of the Saudi Society of Agricultural Sciences* 20: 459–466.

Ding N, Dong H, Ongena M. 2025. Bacterial Cyclic Lipopeptides as Triggers of Plant Immunity and Systemic Resistance Against Pathogens. *Plants* 14: 2644.

Ebrahimi M, Safari Sinegani AA, Sarikhani MR, Aliasgharzad N. 2023. Inoculation effects of isolated plant growth promoting bacteria on wheat yield and grain N content. *Journal of Plant Nutrition* 46: 1407–1420.

Emami S, Alikhani HA, Pourbabaei AA, Etesami H, Sarmadian F, Motessharezadeh B. 2019. Effect of rhizospheric and endophytic bacteria with multiple plant growth promoting traits on wheat growth. *Environmental Science and Pollution Research* 26: 19804–19813.

Farahat MG, Mahmoud M, Youseif SH, Saleh S, Kamel Z. 2020. Alleviation of salinity stress in wheat by ACC deaminase-producing Bacillus aryabhattai EWR29 with multifarious plant growth-promoting attributes. *Plant Archives*.

Franz L, Kazmaier U, W. Truman A, Koehnke J. 2021. Bottromycins - biosynthesis, synthesis and activity. *Natural Product Reports* 38: 1659–1683.

Garrell A-K, Cheadle J, Crook N, Pal G, Septer AN, Wagner MR, Beck AE, Kleiner M. 2026. Differential proteomics of bacteria grown in vitro and in planta reveals functions used during growth on maize roots. *mSystems* 11: e0037126.

Gu Q, Yang Y, Yuan Q, Shi G, Wu L, Lou Z, Huo R, Wu H, Borriss R, Gao X. 2017. Bacillomycin D Produced by Bacillus amyloliquefaciens Is Involved in the Antagonistic

Interaction with the Plant-Pathogenic Fungus Fusarium graminearum. *Applied and Environmental Microbiology* 83: e01075-17.

Hartmann A, Klink S, Rothballer M. 2021. Importance of N-Acyl-Homoserine Lactone-Based Quorum Sensing and Quorum Quenching in Pathogen Control and Plant Growth Promotion. *Pathogens* 10: 1561.

Khourchi S, Elhaissoufi W, Loum M, Ibnyasser A, Haddine M, Ghani R, Barakat A, Zeroual Y, Rchiad Z, Delaplace P, *et al.* 2022. Phosphate solubilizing bacteria can significantly contribute to enhance P availability from polyphosphates and their use efficiency in wheat. *Microbiological Research* 262: 127094.

Kumar R, Singh A, Shukla E, Singh P, Khan A, Singh NK, Srivastava A. 2024. Siderophore of plant growth promoting rhizobacterium origin reduces reactive oxygen species mediated injury in Solanum spp. caused by fungal pathogens. *Journal of Applied Microbiology* 135: lxae036.

Li Y, Li Q, Guan G, Chen S. 2020. Phosphate solubilizing bacteria stimulate wheat rhizosphere and endosphere biological nitrogen fixation by improving phosphorus content. *PeerJ* 8: e9062.

Li Y, Li Y, Zhang H, Wang M, Chen S. 2019. Diazotrophic Paenibacillus beijingensis BJ-18 Provides Nitrogen for Plant and Promotes Plant Growth, Nitrogen Uptake and Metabolism. *Frontiers in Microbiology* 10.

Niu B, Vater J, Rueckert C, Blom J, Lehmann M, Ru J-J, Chen X-H, Wang Q, Borriss R. 2013. Polymyxin P is the active principle in suppressing phytopathogenic Erwinia spp. by the biocontrol rhizobacterium Paenibacillus polymyxa M-1. *BMC Microbiology* 13: 137.

Northen TR, Kleiner M, Torres M, Kovács ÁT, Nicolaisen MH, Krzyżanowska DM, Sharma S, Lund G, Jelsbak L, Baars O, *et al.* 2024. Community standards and future opportunities for synthetic communities in plant–microbiota research. *Nature Microbiology* 9: 2774–2784.

Pang Z, Chen J, Wang T, Gao C, Li Z, Guo L, Xu J, Cheng Y. 2021. Linking Plant Secondary Metabolites and Plant Microbiomes: A Review. *Frontiers in Plant Science* 12.

Qin Z, T. Munnoch J, Devine R, A. Holmes N, F. Seipke R, A. Wilkinson K, Wilkinson B, I. Hutchings M. 2017. Formicamycins, antibacterial polyketides produced by Streptomyces formicae isolated from African Tetraponera plant-ants. *Chemical Science* 8: 3218–3227.

Ravelo-Ortega G, Raya-González J, López-Bucio J. 2023. Compounds from rhizosphere microbes that promote plant growth. *Current Opinion in Plant Biology* 73: 102336.

Selten G, Lamouche F, Gómez-Repollés A, López JL, Zhang X, Smart C, Blahovska Z, Zarate Camargo G, Kelly S, de Jonge R, *et al.* 2026. Functional capacities drive recruitment of bacteria into plant root microbiota. *Nature Microbiology* 11: 2994–3009.

Sharifi R, Sharifi-Tehrani A, Talebi-Jahromi K, Ahmadzadeh M. 2010. Pyoverdine Production in Pseudomonas Fluorescens UTPF5 and its Association with Suppression of Common Bean

Damping off Caused by Rhizoctonia Solani (Kühn). *Journal of Plant Protection Research; 2010; vol. 50; No 1*.

Ullah A, Bano A, Janjua HT. 2020. Chapter 3 - Microbial Secondary Metabolites and Defense of Plant Stress. In: Singh JS, Vimal SR, eds. Microbial Services in Restoration Ecology. Elsevier, 37–46.

Wang Z, Zhang H, Liu L, Li S, Xie J, Xue X, Jiang Y. 2022. Screening of phosphate-solubilizing bacteria and their abilities of phosphorus solubilization and wheat growth promotion. *BMC microbiology* 22: 296.

Zarkani AA, Stein E, Röhrich CR, Schikora M, Evguenieva-Hackenberg E, Degenkolb T, Vilcinskas A, Klug G, Kogel K-H, Schikora A. 2013. Homoserine Lactones Influence the Reaction of Plants to Rhizobia. *International Journal of Molecular Sciences* 14: 17122–17146.

### Supporting Figures

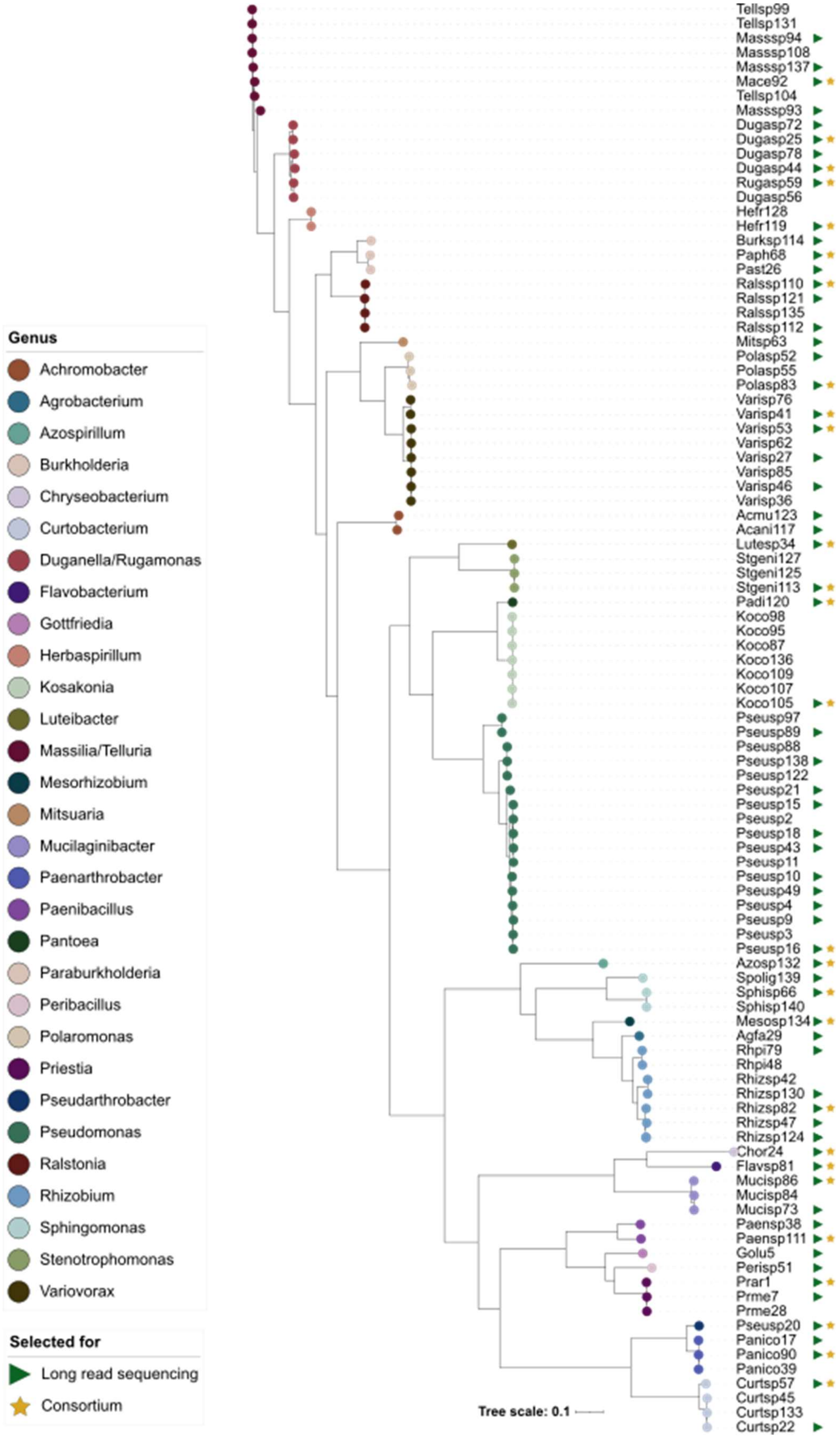
Tellsp99
Tellsp131
Masssp94
Masssp108
Masssp137
Mace92
Tellsp104
Masssp93
Dugasp72
Dugasp25
Dugasp78
Dugasp44
Rugasp59
Dugasp56
Hefr128
Hefr119
Burksp114
Paph68
Past26
Ralssp110
Ralssp121
Ralssp135
Ralssp112
Mitsp63
Polasp52
Polasp55
Polasp83
Varisp76
Varisp41
Varisp53
Varisp62
Varisp27
Varisp85
Varisp46
Varisp36
Acmu123
Acani117
Lutesp34
Stgeni127
Stgeni125
Stgeni113
Padi120
Koco98
Koco95
Koco87
Koco136
Koco109
Koco107
Koco105
Pseusp97
Pseusp89
Pseusp88
Pseusp138
Pseusp122
Pseusp21
Pseusp15
Pseusp2
Pseusp18
Pseusp43
Pseusp11
Pseusp10
Pseusp49
Pseusp4
Pseusp9
Pseusp3
Pseusp16
Azosp132
Spolig139
Sphisp66
Sphisp140
Mesosp134
Agfa29
Rhpi79
Rhpi48
Rhizsp42
Rhizsp130
Rhizsp82
Rhizsp47
Rhizsp124
Chor24
Flavsp81
Mucisp86
Mucisp84
Mucisp73
Paensp38
Paensp111
Golu5
Perisp51
Prar1
Prme7
Prme28
Pseusp20
Panico17
Panico90
Panico39
Curtsp57
Curtsp45
Curtsp133
Curtsp22
Tree scale: 0.1
Genus
Achromobacter
Agrobacterium
Azospirillum
Burkholderia
Chryseobacterium
Curtobacterium
Duganella/Rugamonas
Flavobacterium
Gottfriedia
Herbaspirillum
Kosakonia
Luteibacter
Massilia/Telluria
Mesorhizobium
Mitsuaria
Mucilaginibacter
Paenarthrobacter
Paenibacillus
Pantoea
Paraburkholderia
Peribacillus
Polaromonas
Priestia
Pseudarthrobacter
Pseudomonas
Ralstonia
Rhizobium
Sphingomonas
Stenotrophomonas
Variovorax
Selected for
Long read sequencing
Consortium

222

Fig S1. Phylogenetic tree of 99 sequenced isolates based on 16S rRNA gene sequences. The sequence alignment and tree were produced using the SILVA Alignment, Classification, and Tree (ACT) online tool. The tree was inferred using FastTree under the GTR + Gamma model, with branch lengths representing the number of substitutions per site. Green arrows indicate strains selected for long-read sequencing. Strains marked by a yellow star indicate consortium membership.

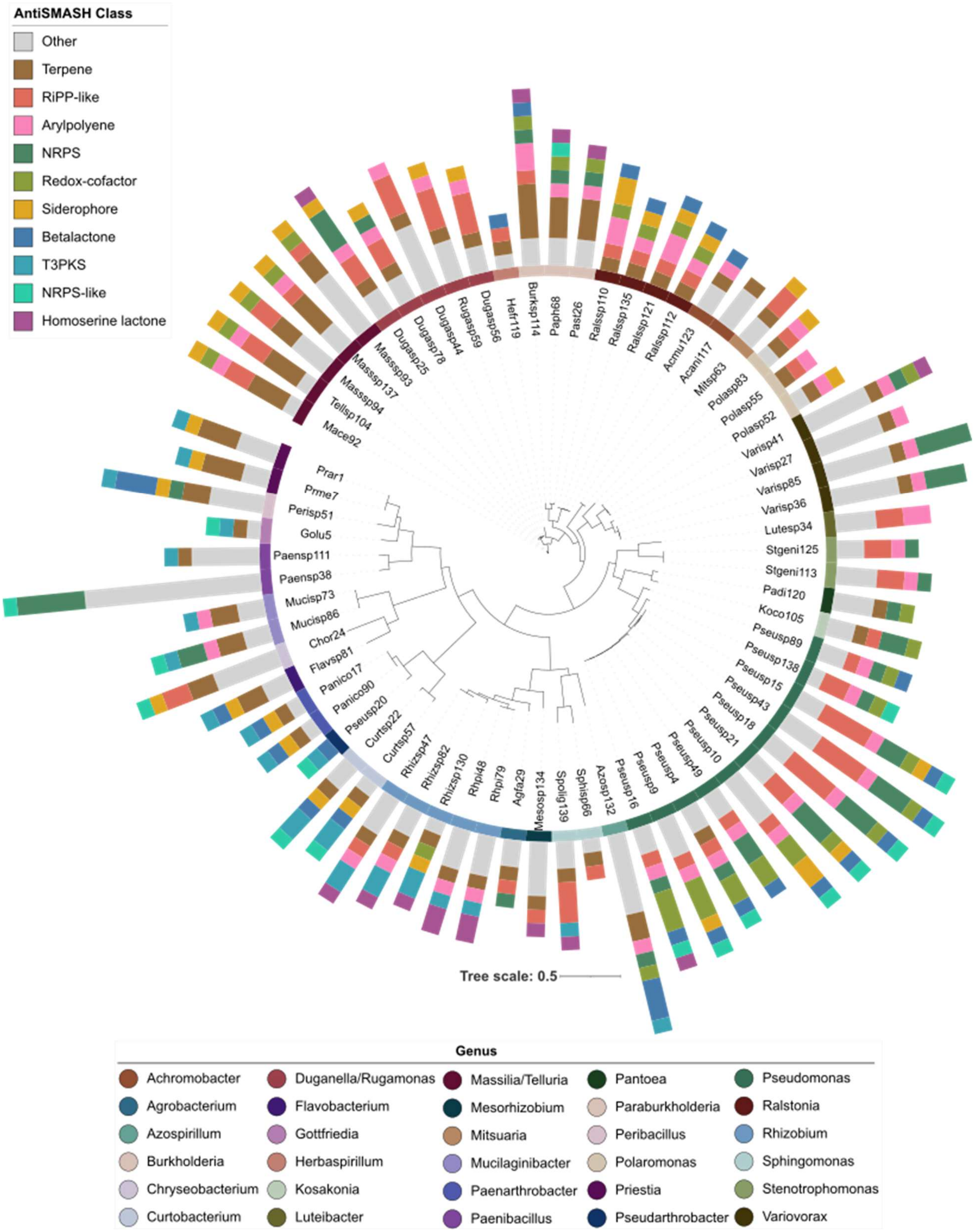


Fig S2. Distribution of the top ten biosynthetic gene clusters (BGCs) identified across representative isolates (n=69) from each clade in the TriMic collection available at the DSMZ. Stacked bar plots indicate the number of BGCs identified for each AntiSMASH class. The top ten identified BGCs classes include: Terpenes, Aryl polyene (arylpolyene), non-ribosomal peptide synthetase (NRPS), redox-cofactors such as PQQ (Redox cofactor), siderophores, beta-lactone containing protease inhibitor (Betalactone), type III polyketide synthase (T3PKS), NRPS-like fragment (NRPS-like), and homoserine lactone.

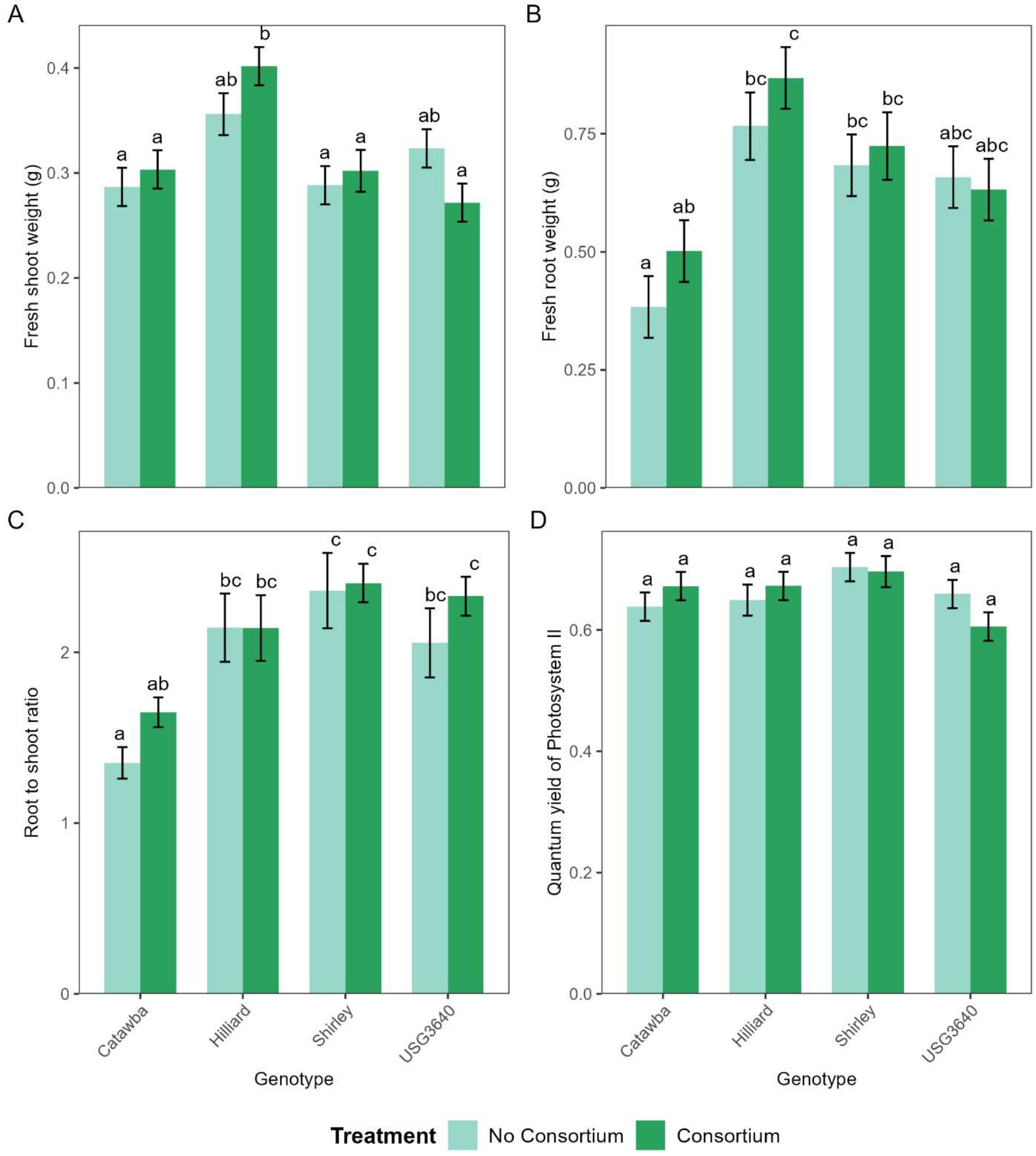


Fig S3. Mean wheat responses to consortium inoculation at 15 days post emergence (Mean ± standard error). A) Fresh shoot weight, B) Fresh root weight, C) Root to shoot ratio, and D) Photosystem II quantum yield (chlorophyll fluorescence). Different letters indication significant differences based on Tukey- adjusted pairwise comparisons at α= 0.05.

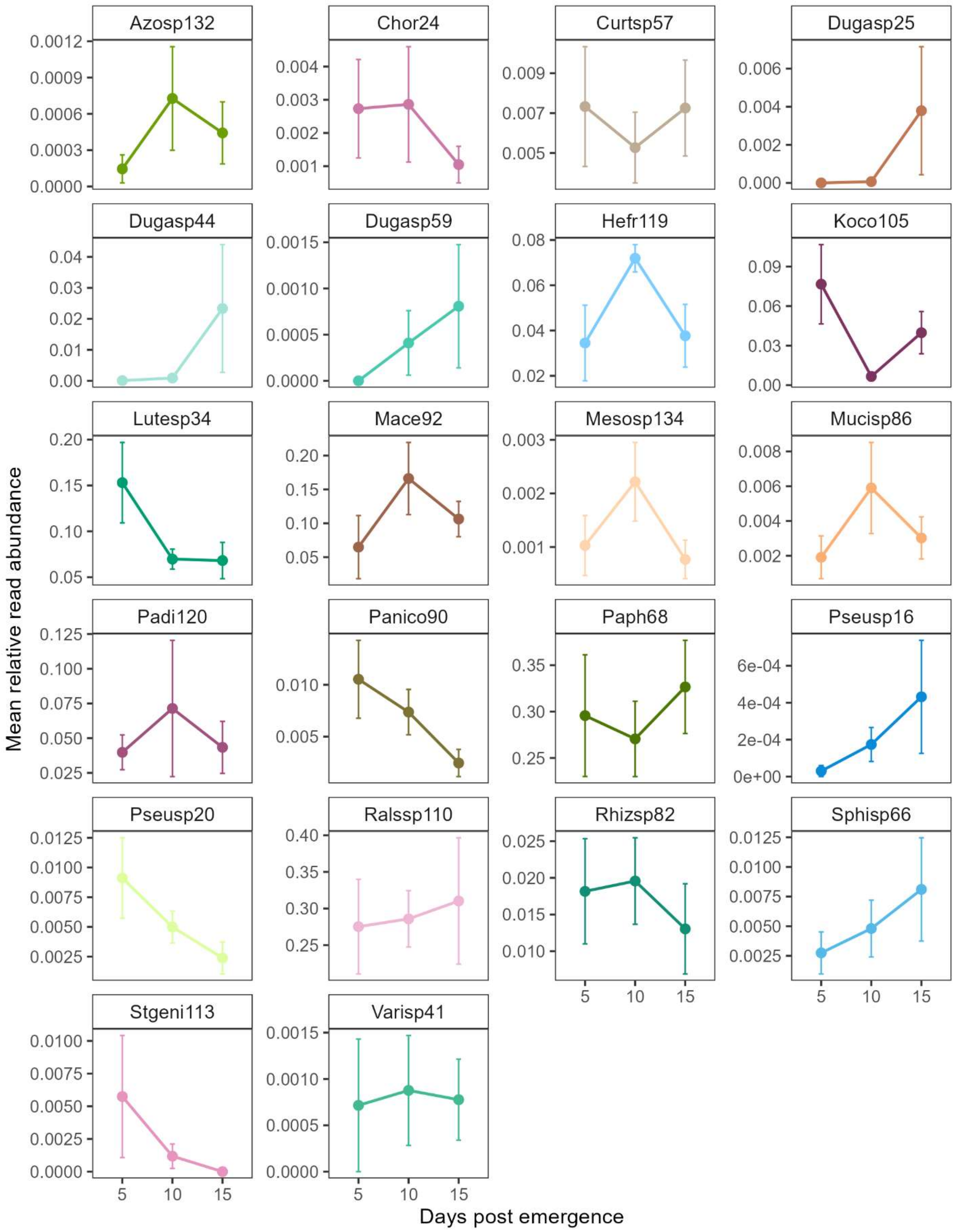


Fig S4. Relative read abundance (mean±SE) of individual consortium strains in the wheat cultivar Catawba at 5, 10, and 15 days post-emergence (DPE). Only consortium strains with relative read abundances >0.01% are shown.

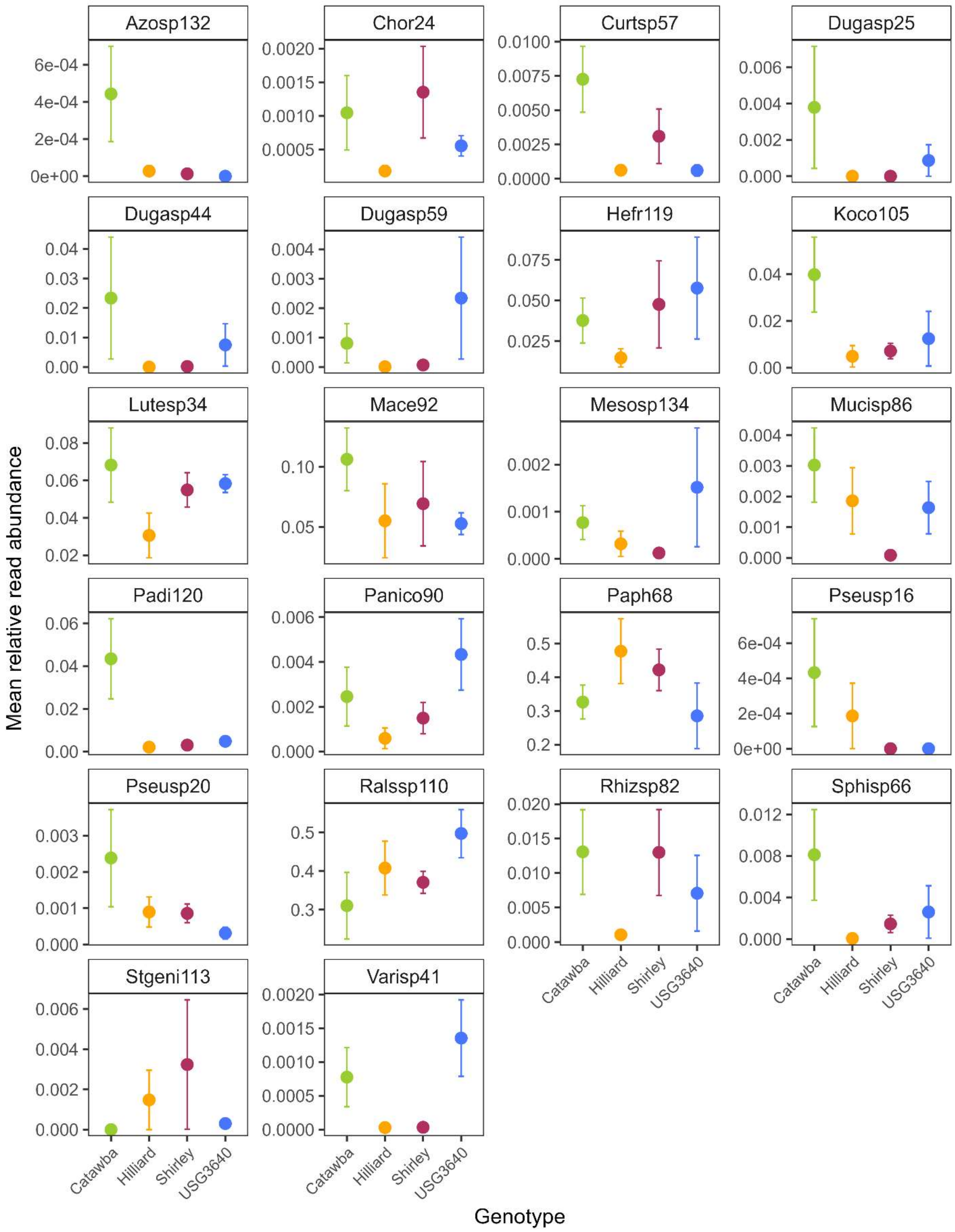


Fig S5. Relative read abundance (mean±SE) of individual consortium strains in across the four wheat genotypes at 15 days post-emergence (DPE). Only consortium strains with relative read abundances >0.01% are shown.

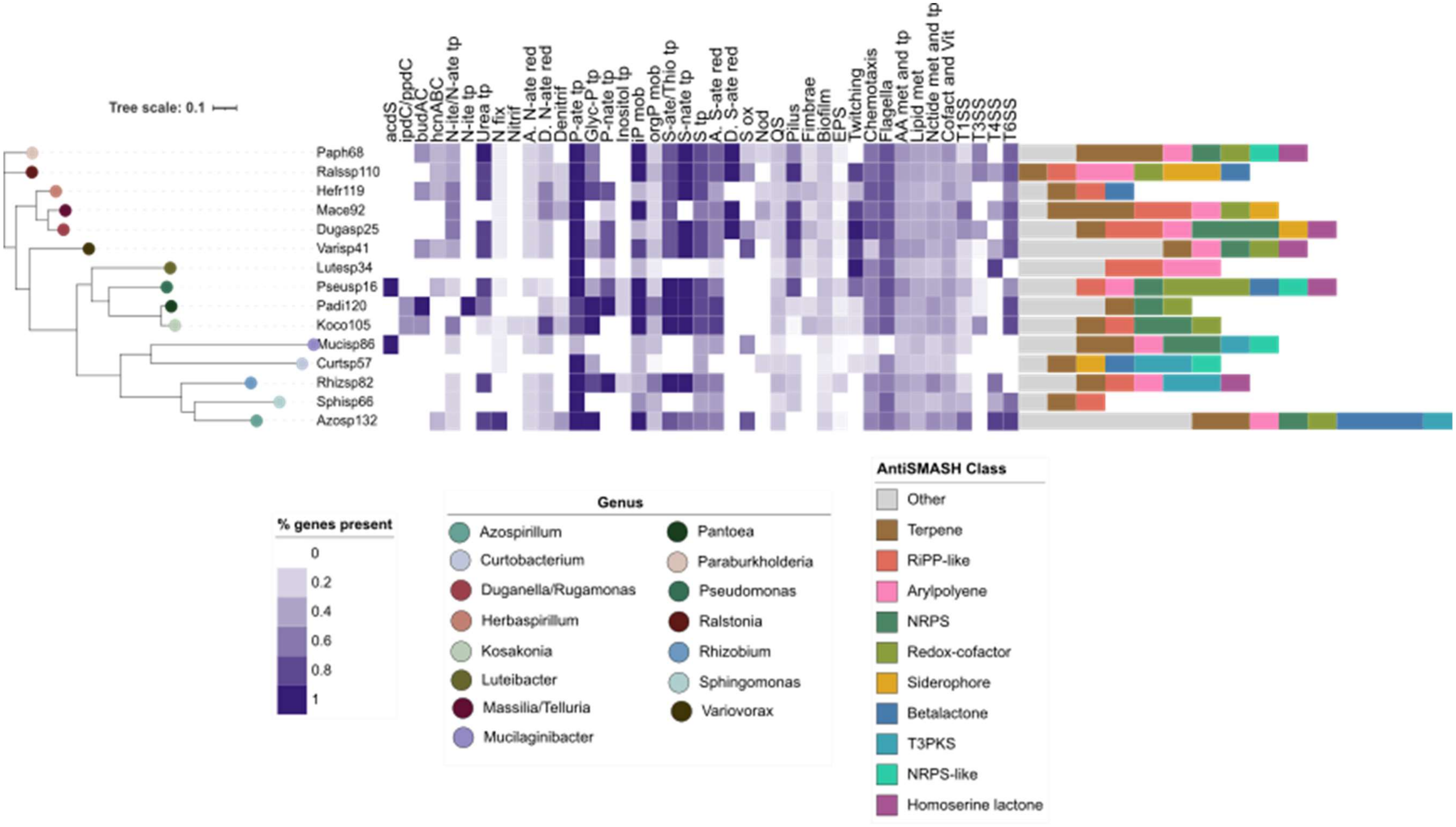


Fig S6. Wheat root synthetic community (SynCom) functional traits. The heatmap depicts the proportion of genes detected per genome within the selected functional categories, calculated as the number of genes present relative to the total number of genes in the category. Functional categories (n= number of genes in category) are as follows: Plant growth promotion: 1-aminocyclopropane-1-carboxylate deaminase (acdS, n = 2), indole-3-acetic acid synthesis (ipdC/ppdC, n = 2), acetoin/2,3-butanediol synthesis (budAC, n = 2), and hydrogen cyanide production (hcnABC, n = 5). Nitrogen and phosphorus cycling: nitrate/nitrite transport (N-ite/N-ate tp, n = 5), nitrite transport (N-ite tp, n = 1), urea transport (Urea tp, n = 6), nitrogen fixation (N fix, n = 11), nitrification (Nitrif, n = 5), assimilatory nitrate reduction (A. N-ate red, n = 5), dissimilatory nitrate reduction (D. N-ate red, n = 7), denitrification (Denitrif, n = 13), phosphate transport (P-ate tp, n = 6), glycerol-phosphate transport (Glyc-P tp, n=5), phosphonate transport (P-nate tp, n = 4), inositol transport (Inositol tp, n = 9), inorganic phosphorus mobilization (iP mob, n = 6), and organic phosphorus mobilization (orgP mob, n = 21). Adaptation to plants and colonization: sulfate/thiosulfate transport (S-ate/Thio tp, n=4), sulfonate transport (S-nate tp, n = 3), sulfur relay system (SRS, n = 5), assimilatory sulfate reduction (A. S-ate red, n = 7), dissimilatory sulfate reduction (D. S-ate red, n = 1), sulfur oxidation (S ox, n = 4), nodulation (Nod, n = 4), quorum sensing (QS, n = 68), pilus formation (Pilus, n = 8), fimbriae (Fimbriae, n = 6), biofilm formation (Biofilm, n = 85), exopolysaccharide biosynthesis (EPS, n = 36), twitching motility (Twitching, n=6), chemotaxis (Chemotaxis, n = 27), flagellar motility (Flagella, n = 42), amino acid metabolism and transport (AA met and tp, n=451), lipid metabolism (Lipid met, n = 107), nucleotide metabolism and transport (Nctide met and tp), metabolism of cofactors and vitamins (Cofact and Vit), and secretion systems including Type I (T1SS, n = 6), Type III (T3SS, n = 24), Type IV (T4SS, n = 19), and Type VI (T6SS, n = 20). Stacked bar plots indicate the number of BGCs identified for each AntiSMASH class. The top ten identified BGCs classes include: Terpenes, Aryl polyene (arylpolyene), non-ribosomal peptide synthetase (NRPS), redox-cofactors such as PQQ (Redox cofactor), siderophores, beta-lactone containing protease inhibitor (Betalactone), type III polyketide synthase (T3PKS), NRPS-like fragment (NRPS-like), and homoserine lactone.
